# Synergic coordination of stem cells is required to induce a regenerative response in anthozoan cnidarians

**DOI:** 10.1101/2019.12.31.891804

**Authors:** Aldine R. Amiel, Kevin Foucher, Solène Ferreira, Eric Röttinger

## Abstract

Little is known about the origin of the inductive signal that translates the amputation stress into a cooperative cellular response. By studying the process underlying the reformation of lost body parts in the anthozoan cnidarian *Nematostella vectensis*, we identified a regeneration-inducing structure that, via a tissue crosstalk, is responsible for the initiation of the repair program. We further revealed for the first time in anthozoan cnidarians, that fast and slow-cycling/quiescent stem cells respond to the amputation stress and actively participate in the reformation of lost body parts. Importantly, a synergic interaction of both stem cell populations is required to complete the regeneration process. Our findings suggest that the emergence/loss of structure complexity/compartmentalization influences the proprieties of tissue plasticity, changes the competence of a tissue to reprogram and, in the context of regeneration, the capacity of the tissue to emit or respond to a regeneration-inducing signal.

## Background

The process of regeneration is the re-formation of damaged or missing body parts that in bilaterian animals, involves cellular migration and proliferation of adult stem cells or de-differentiated cell populations. In combination with the remodeling of existing cells/tissues, these cellular processes are crucial to re-establish the symmetry and the proportion of the body [1]. Regeneration is widespread among metazoans and has fascinated scientists for centuries [2, 3]. Still, our understanding of why some animals regenerate while others can’t remains largely unknown. In the past 10 years, historical invertebrate whole-body regeneration models such as Planaria and Cnidaria have re-emerged and are taking advantage of modern cellular and molecular techniques to understand the complex process of regeneration [4–7]. Important insight has been gained from these species that have increased our understanding about the cellular and molecular mechanisms underlying regeneration. Yet little is known about how the organism translates the amputation stress into a regeneration inducing signal and the origin as well as the nature of the inductive signal required for the cooperation of cell populations during the regeneration process.

The level of regenerative abilities varies across the animal kingdom [7]. While vertebrates can regenerate certain structures and tissues, planarians and cnidarians posses extraordinary whole-body regeneration capacities and can regrow missing body parts when dissected into small pieces [2, 8]. Nonetheless, their regenerative capacities have limitation at both the species and also the tissues level. For instance in planarians, isolation of the most anterior part of its body (region in front of the photoreceptor) or the pharynx does not lead to regeneration as these regions are devoid of planarian specific pluripotent stem cells, the neoblasts [9–11]. In *Hydra*, the isolated body column, named gastric region, possesses the ability to regenerate while the isolated head, pedal disk or tentacles cannot. The absence of regeneration from these isolated parts is due to the lack of hydrozoan specific pluripotent stem cells, the i-cells (interstitial stem cells), in these highly differentiated and specialized regions of the body [12, 13]. Thus, identifying the regenerative limits of a given organism and using them for experimental approaches has led to important findings on the location of stem cell populations and their dynamics during the complex biological process of regeneration.

To date, the majority of studies addressing the role of stem cells during regeneration in cnidarians were performed using Hydrozoa such as *Hydra* and *Hydractinia* [2,14–17]. While *Hydra* is well known to have developed an alternative mechanism to regenerate a head in a stem-cell (i-cell) depleted context [18], those classically considered fast cycling i-cells are under “normal” conditions involved in the regeneration process in *Hydra* and *Hydractinia* [1,19–23]). Only recently, a study in *Hydra* has described slow cycling cells that are part of the pluripotent i-cell as well as the unipotent epithelial stem cell populations and involved in regeneration [24]. However, nothing is known about the relationship between fast and slow cycling stem cells, their hierarchy and how they are interacting to trigger tissue reprograming for regeneration. This question remains unanswered not only in hydrozoans, but in cnidarians in general.

Cnidarians are composed of two major groups, the Anthozoa (coral, sea anemone) and the Medusozoa, that include Hydrozoa, Schyphozoa and Cubozoa [25–30]. Interestingly, the crown-group cnidarians (the living representatives) are genetically as divergent as protostomes and deuterostomes [31, 32] and it has been proposed that none of the cnidarian clades is representing the entire group [32]. Thus, it is of great interest to compare the regenerative strategies between hydrozoan and non-hydrozoan cnidarians in order to understand the similarities/differences and evolution of the cellular and molecular mechanisms driving regeneration. Importantly, the presence of adult stem cells has yet to be discovered in non-hydrozoan cnidarians.

Anthozoa diverged early within the cnidarian phylogeny [29, 33] and form the largest group of cnidarian [34, 35] that display anatomical differences compared to other cnidarian polyps. Exclusively anthozoans possess mesenteries that are well-defined anatomical structures located in the gastric cavity, fulfilling the digestive and sexual reproduction roles of the individual polyp. Mesenteries are multifunctional tissues of combined ectodermal and endodermal origins that in addition to the germ lineage and digestive cells also possesses various additional cell types such as muscle cells and cnidocytes (stinging cells) [36–38]. Because of its i) phylogenetic position, the ii) ease to control spawning and fertilization [39–41] and thus the access to embryonic, larval and post-metamorphic stages all year round, iii) a sequenced genome [42] as well as iv) available cellular and molecular tools for functional genomics [43, 44] and genome editing [45], the anthozoan sea anemone *Nematostella vectensis* has become a leading model to gain new insight into animal evolution [42,46,47] and early cnidarian development [48–55]. *Nematostella* possesses also high regenerative abilities and therefore is a potent model to better understand the relationship between embryonic development and regeneration [44, 56]. However, our knowledge concerning the morphological, cellular and molecular mechanisms underlying whole body regeneration in *Nematostella* is still in its early phase [57–61]. Recent work has begun to describe the morphological and cellular events involved in the regeneration process in *Nematostella* [59,60,62]. Wound healing after head amputation is already completed within six hours post amputation (hpa) and oral regeneration following sub-pharyngeal amputation occurs in stereotypic steps [62]. This process is first visible when the most oral part of the remaining mesenteries fuses together and enters in contact with the epithelia at the amputation site. The most oral part of the mesenteries is at the origin of the internal part of the pharynx that completes its regeneration at the same time than the tentacles bulbs form and elongate [62]. The various stages of the polyp’s life are defined by clear morphological characteristics (i.e. number of tentacles and mesenteries, presence of gametes) and cell proliferation is necessary for oral regeneration in juveniles [59] as well as in adults [62]. Nonetheless, in juvenile as well as in adult *Nematostella*, the first morphogenetic step of regeneration is proliferation independent and only later steps require mitotic activity to fully complete oral regeneration following sub-pharyngeal amputation [62]. While this study laid down a foundation for functional work on anthozoan regeneration, basic but crucial questions about the regenerative capacities in *Nematostella* still remain unanswered: i) are there any body part, tissue and/or age specific limits of its regenerative abilities? ii) what is the cellular/tissue origin of the signal(s) that initiates cell proliferation and/or regeneration? iii) what are the cellular mechanisms underlying this phenomenon, and importantly, iv) are there adult stem cells in anthozoans and v) are they involved in the regeneration process?

In the present study we combined classical grafting experiments with cellular approaches to address these questions. This strategy enabled us to highlight a crucial crosstalk between the mesenteries and the surrounding epithelia of the body wall required for regeneration and to identify that the mesenteries are crucial to induce proliferation/regeneration in *Nematostella*. We further identified at least two cell populations, a fast cycling one and a quiescent/slow cycling one that are required in a combinatorial manner to trigger a regenerative response. Both of these cell populations are re-activated in response to the amputation stress and participate in the regeneration process. Importantly, our work provides the first set of evidences for the existence of at least two adult stem cell populations in anthozoans that synergize to drive efficient regeneration in the anthozoan cnidarian *Nematostella*.

## Materials and methods

### Animal culture

*Nematostella vectensis* were cultivated at the Institute for Research on Cancer and Aging, Nice (IRCAN) of the University of Nice. Adult and juvenile animals were cultured in 1/3X ASW (Artificial seawater, Tropic-Marin Bio-actif system sea salt) and maintained at 17°C or 22°C for adults and juveniles, respectively. Adults were fed five times a week with freshly hatched artemia. Spawning to obtain juveniles was carried out as described in [39]. Juveniles for cutting experiments were raised until six weeks after fertilization. Starting 2 weeks post fertilization, they were feed once a week with 1ml of smashed artemia during four weeks.

### Cutting, tissue isolation and grafting experiments

Juveniles or adults were relaxed by adding 1 ml of 7.14% MgCl_2_ in 5ml 1/3 ASW and left on a light table during 10 to 15 min until they were relaxed enough for surgical interventions. Polyps were cut using a microsurgery scalpel (Swann-Morton, n°15). Each cut was performed perpendicular to the oral-aboral axis of the body column. Grafting experiments were performed using adult tissues only from a clonal colony, following the cutting protocol described above and carried as follows: 1) Head and physa were surgically removed from relaxed animals; 2) The body column was isolated and opened along the oral-aboral axis of the polyp with microsurgical scissors (Fine Science Tools GMBH #15100-09, Heidelberg, Germany); 3) If required, the mesenteries were gently removed mechanically from the epithelia using forceps (Rubis Grip Tweezers, style 3C) and microsurgical scissors 4) Isolated epithelia from the body wall were forced to contact the freshly isolated mesenteries approximately 5 to 10 min after tissue isolation and the graft was maintained with a human hair allowing the wounds to heal properly; 5) After three days the hair was removed to avoid any mechanical constraints or tissue damages during the regenerating period. Isolated fragments and grafts were transferred into a new dish (or rinsed at least five times to remove MgCl_2_) and maintained in 1/3 ASW at 22°C (if not stated otherwise) for the duration of the experiments.

### Fixation, DNA, actin labeling

After relaxing adult or juvenile polyps as well as isolated body parts in MgCl_2_ for 10-15 minutes, animals were fixed in 4% paraformaldehyde (Electron Microscopy Sciences # 15714) in 1/3 ASW during 1 hour at 22°C or overnight at 4°C. Fixed animals were washed three times in PBT 0.2% (PBS1x + Triton 0.2%). To analyze the general morphology, Hoechst staining (Invitrogen #33342, Carlsbad, CA, USA) at 1/5000 was used to label the DNA/nucleus and BODIPY® FL PhallAcidin 488 (Molecular Probes #B607, Eugene, OR, USA) staining at 1/200 to label actin microfilament (cell membranes and muscle fibers).

### Cell proliferation and “pulse and chase” experiment

To detect cellular proliferation Click-it EdU (5-ethynyl-2’-deoxyuridine) kits (Invitrogen # C10337 or # C10339, Carlsbad, CA, USA) were used following the protocol from [59]. To visualize cell proliferation at a given time point EdU incubation was carried out during 30min. For the “pulse and chase” experiment EdU incubation was performed during 1h, then washed 5 times and chased for the period of interest. In order to identify of the Label Retaining Cells (LRCs), animals were incubated for one week with EdU, then chased up to 11 weeks for juveniles or 3 to 5 month for adults.

### Irradiation

For irradiations experiments, animals where placed in a 10cm diameter petri dish with 15ml 1/3 ASW. Irradiations at 100 and 300 Grays where performed on the juvenile polyps using a X-ray irradiator (CP-160 Cabinet X-Radiator, Faixitron). After irradiation, animals were recovering during 4 hours in their culture medium (1/3ASW) at room temperature before further cutting experiments.

### Imaging

Live animals were imaged using a protocol described in [63]. The imaging setup was composed of either with a Zeiss Stereo Discovery V8 Discovery or a Zeiss Axio Imager A2 (both Carl Zeiss Microscopy GmbH, Jena, Germany) equipped with a Canon 6D (Canon) or a Axiocam 506 color camera (Zeiss) digital camera, triggering two external Canon Speedlite 430 EX II Flashes and controlled by the Canon Digital Photo Professional software (Canon Inc, Tokyo, Japan). Images were edited using Adobe Lightroom 5 and/or Photoshop CS6 software (Adobe Systems Inc, San Jose, CA, USA). Labeled animals were analyzed using a Zeiss LSM Exciter confocal microscope running the ZEN 2009 software (Carl Zeiss Microscopy GmbH, Jena, Germany) from the IRCAN imaging platform (PICMI). Each final image was reconstituted from a stack of confocal images using Z-projection (maximun intensity or standard deviation) of the ImageJ software (Rasband, W.S., ImageJ, U. S. National Institutes of Health, Bethesda, Maryland, USA, http://imagej.nih.gov/ij/, 1997-2014).

## Results

### The regenerative capacity of the physa is age dependent

In order to assess if all body parts of *Nematostella* possess similar regenerative capacities and if those limits are age dependent, we dissected adult and juvenile *Nematostella* into several isolated body parts. Following dissection, we analyzed each parts capacity to regenerate the missing oral or aboral structures within seven days (Figures 1, Additional file 1: Figure S1). Throughout the manuscript the isolated body parts that are tracked to score regeneration are indicated as follow: [name of the isolated part]. As an example, when a tentacle is isolated and tracked for it is ability to regenerate, it is indicated as [tentacle]. [full body column + physa] is the equivalent of the sub-pharyngeal amputation experiment performed in previous studies [59, 62] (Figures 1Ac,c’, Bh,h’, C) and is considered as positive control in the corresponding experiments.

**Figure 1.**
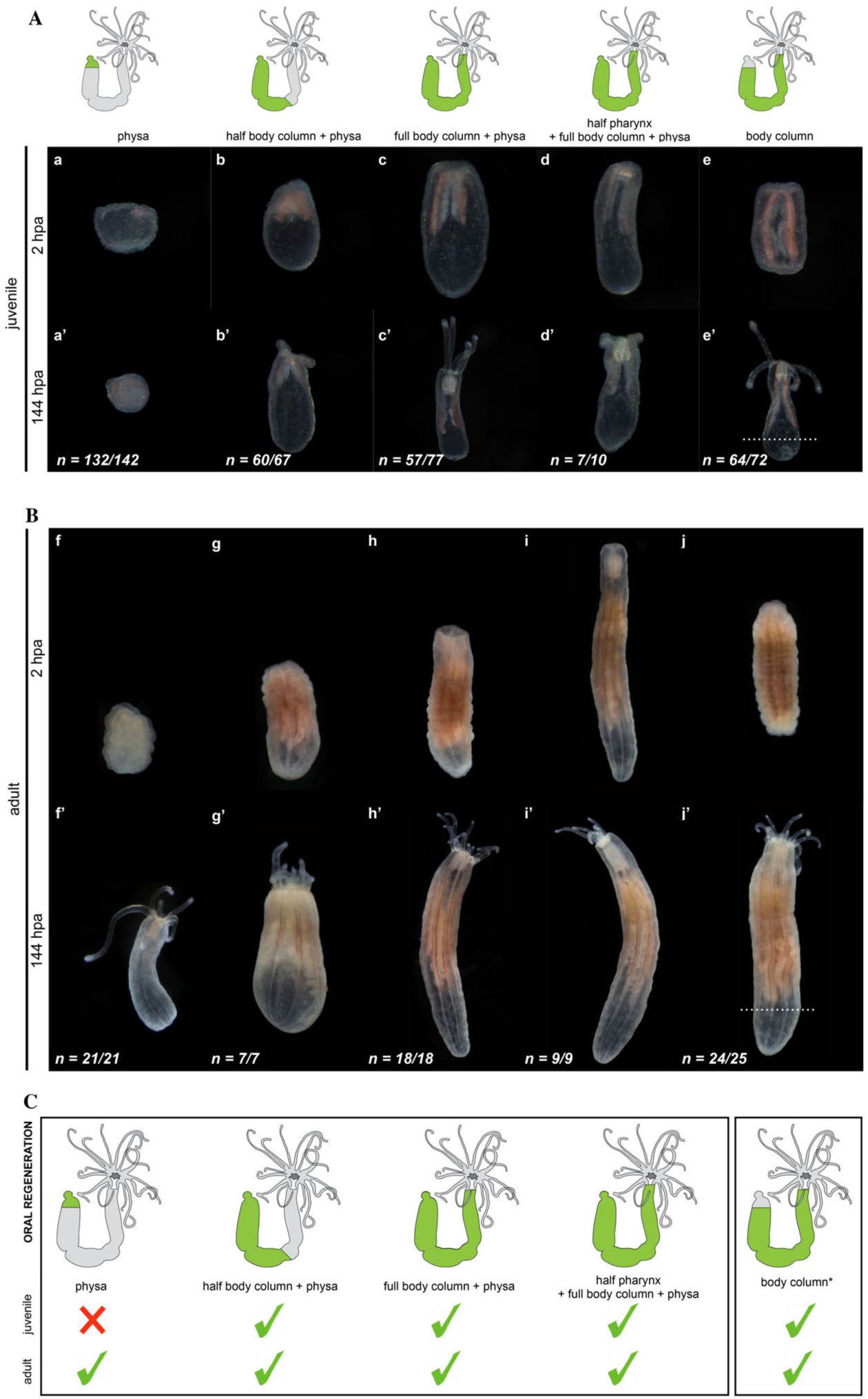
Oral regenerative capacity analyzed in juveniles (Aa-e, Aa’-e’) and adults (Bf-j, Bf’-j’) from isolated [physa] (Aa, Aa’, Bf, Bf’); [half body column + physa] (Ab, Ab’, Bg, Bg’); [full body column + physa] (Ac, Ac’, Bh, Bh’); [half pharynx + full body column + physa] (Ad, Ad’, Bi, Bi’); [body column] (Ae, Ae’, Bj, Bj’) from which aboral regeneration was also assessed (dashed line in Ae’, Bj’). On top of each panel are *Nematostella* illustrations in which each isolated part is indicated in green and the part scored for regeneration in grey. Photographs of the isolated body parts at 2hpa (Aa-e, Bf-j). Phenotypes observed after 144hpa (6 days post amputation) (Aa’-e’, Bf’-j’). n=[number of specimen with represented phenotype]/[total number of analyzed specimen]. **C**. Diagram summarizing the oral regeneration experiments carried out in juveniles and adults. Green parts in the schematic *Nematostella* indicate the isolated body part, green checkmarks regenerative success and red crosses the absence of regeneration after body part isolation.

We first addressed the capacity of isolated body parts to regenerate missing oral structures (oral regeneration) and we observed that most body regions isolated from either juvenile or adult polyps regenerate within 144hpa. The only exception was the isolated juvenile (but not adult) physa that failed to regenerate missing oral features (pharynx, mouth, tentacles, Figure 1Aa-e, a’-e’, Bf-j, f’-j’, C). The capacity of body parts to regenerate is correlated with the presence of massive cell proliferation at the amputation site throughout regeneration (Figure 2A, B). Massive cell proliferation was also observed at the amputation site (Figure 3Aa,b) and is required for regeneration of the adult [physa] (Figure 3Ac,d). Interestingly, no or very little cell division was detected in juvenile [physa] (Figure 3Ba,b). These observations show that for the various isolated body parts the capacity to regenerate is associated to cell proliferation, suggesting similar cellular mechanisms involved in this process along the different regions of the polyp. They also highlight that existing age-dependent differences in the regenerative capacity of *Nematostella* are body part specific.

**Figure 2.**
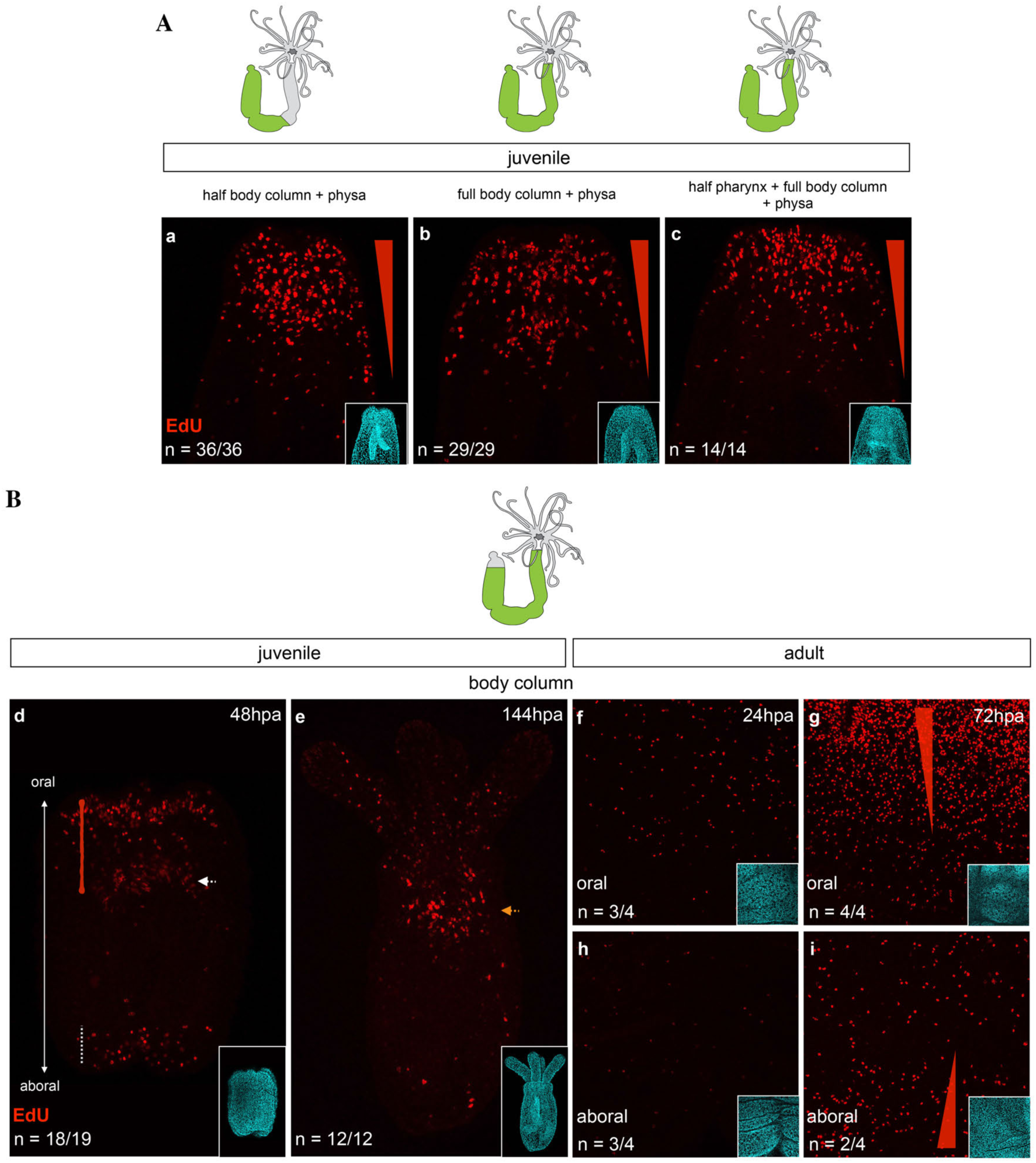
**A.** Cell proliferation in juveniles at 48hpa for [half body column + physa] (a), [full body column + physa]](b) and [half pharynx + full body column + physa] (c). **B**. Cell proliferation in juveniles at 48 (d) and 144hpa (e) and in adult at 24 (f, oral; h, aboral) and 72hpa (g, oral; i, aboral) for [body column]. The left red triangle in g and i represent a gradient of cell proliferation. Confocal images of cell proliferation in red (EdU) (Aa-c, Bd-i) and DNA/nucleus in cyan (DAPI) (small insert in Aa-c, Bd-i). n=[number of specimen with represented phenotype]/[total number of analyzed specimen].

**Figure 3.**
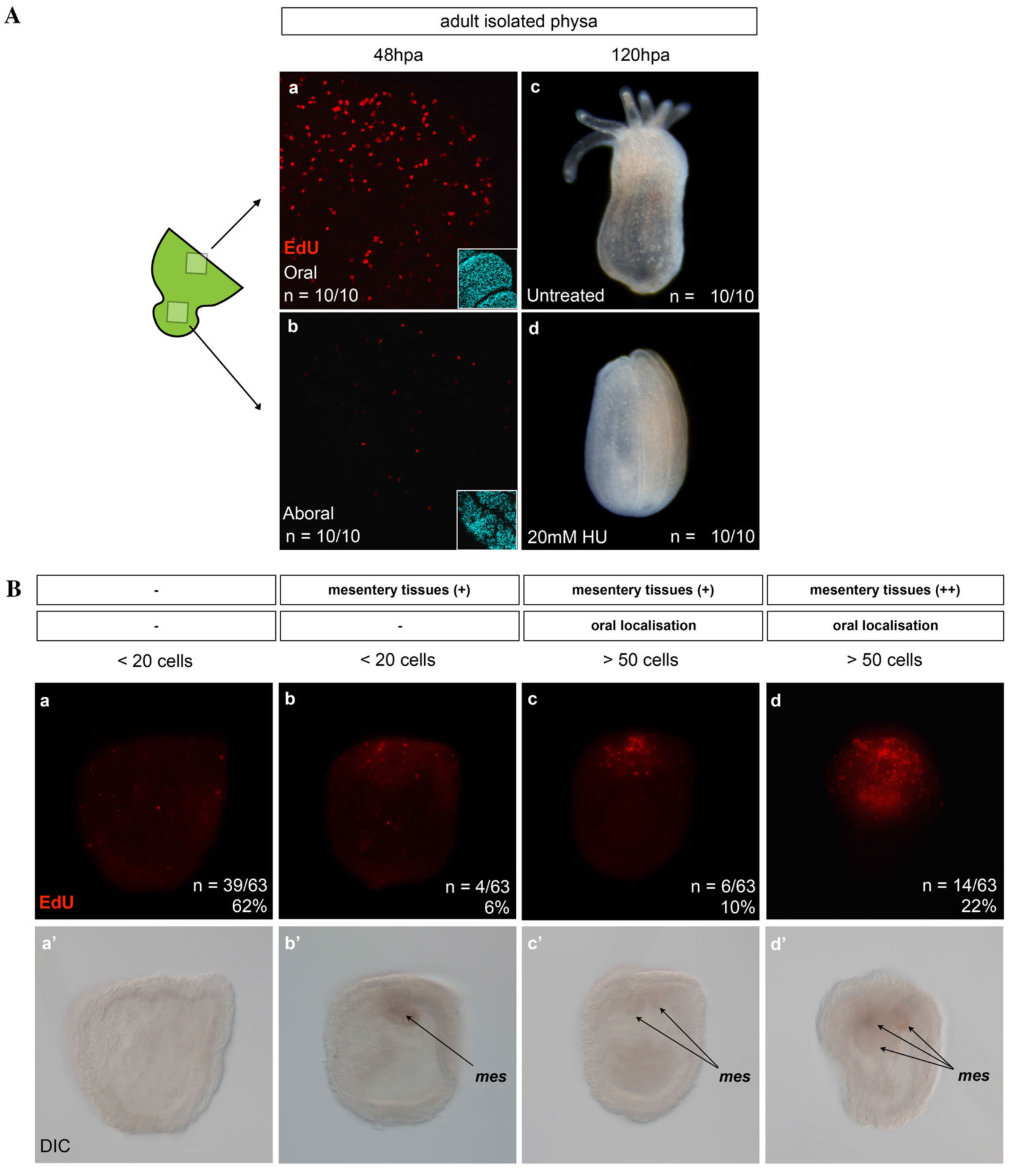
Cell proliferation in adult (**A**) and juvenile (**B**) [physa]. **A**. In adults, cell proliferation is present at the amputation site (Aa,b) and required for regeneration (c,d) of the [physa]. (a,b) Confocal microscopy images showing cellular proliferation in red (EdU) at 48hpa (a,b). Oral most part (a) and aboral most part (b) of the adult isolated physa. The cartoon to the left indicates the isolated physa and the regions represented in (a) and (b). (c,d) Hydroxyurea (HU) at 20mM was used in this set of experiment to inhibit cell proliferation in adult [physa] (untreated control in c, HU treated animal in d). n=[number of specimen with represented phenotype]/[total number of analyzed specimen]. **B**. Cell proliferation is absent/reduced in isolated juvenile physa at 48hpa. Observation of cellular proliferation at 48hpa using EdU staining on isolated juvenile physa. (a-d) Fluorescence microscopy images showing cellular proliferation (green), (a’-d’) corresponding DIC images of the isolated physa represented above. Black arrows indicate the position of remaining mesentery tissue. On top of the panel the number of EdU positive cells (< 20 cells to > 50 cells), the amount of mesentery tissue (-, absent, +, few or ++, more than few) as well as the localization of EdU staining (-, random, oral) within the isolated physa are represented. n=[number of specimen with represented phenotype]/[total number of analyzed specimen]

A similar conclusion was obtained when analyzing the regenerative capacity of isolated juvenile or adult oral body regions to reform aboral structures (aboral regeneration) (Additional file 2: Figure S2). Only juvenile [tentacles + full pharynx + half body column] and [body column], as well as adult [tentacles + full pharynx], [tentacles + full pharynx + half body column] and [pharynx] regenerated (Additional file 1: Figure *S2Ad, d’, Bg-j, g’-j’, C*). Intriguingly though, we did not observed massive cell proliferation at the amputation site, neither for the regenerating nor the non-regenerating isolated body parts (Additional file 3: Figure S3), suggesting different cellular strategies involved in the oral *vs* aboral regeneration processes.

Taken together these experiments clearly illustrate the existence of limits in the regenerative abilities of distinct *Nematostella* body parts (i.e. the tentacles) that is also varying in an age-dependent manner (i.e. physa). While aboral regeneration seems to involve mainly cell/tissue rearrangements, importantly, the capacity of body parts to reform missing oral structures is tightly linked to the ability of cells to re-enter mitosis, suggesting qualitative variations in the tissues that form the body of *Nematostella*.

### Mesenteries are present in the adult but not in the juvenile physa

Intrigued by the observation that some body parts were able to perform oral regeneration when isolated from adults but not from juveniles (e.g. physa, Figure 1Aa,a’, Bf,f’, C), we analyzed in detail the anatomy of the physa in juvenile *vs* adult polyps (Figure 4). DIC microscopy revealed that in addition to the size variation between juvenile and adult physa, a major difference exists in the thickness and elongation of the longitudinal muscle fiber extensions within the mesenteries (Figure 4a,c). While in adult physa we observed opaque longitudinal extensions from the aboral end of the mesenteries towards the aboral-most tips of the physa (Figure 4c), in juvenile physa, we couldn’t detect those longitudinal extensions (Figure 4a). We further performed confocal imaging of the juvenile and adult [physa] using phalloidin to label actin microfilaments (longitudinal and transversal muscle fibers as well as cell cortex) and DAPI to label the nucleus (Figure 4b,b’,d,d’). We observed that in adult [physa], the longitudinal muscle fibers are considerably thicker compared to the ones present in juvenile tissues (Figure 4b,d). Orthogonal projection of these confocal images further revealed significant differences in the organization of the longitudinal muscle fibers in juvenile or adult [physa] epithelia (Figure 4b’,d’). In juvenile [physa], we observed a discrete protrusion from the longitudinal muscle fibers toward the gastric cavity (Figure 4b’). In contrast, the longitudinal muscle fibers of the adult [physa] are thick and shaped in a characteristic manner (Figure 4d’). Within the adult endoderm, they form a stalk at the base of the epithelium that becomes undulated more distally (Figure 4d’). Interestingly, this particular form is strikingly similar to the organization of parietal muscles that support the developing mesenteries [36]. Thus, these observations show the presence of forming mesenteries only in the adult physa and suggest a correlation between the presence of the mesenteries and the capacity of a given tissue to regenerate.

**Figure 4.**
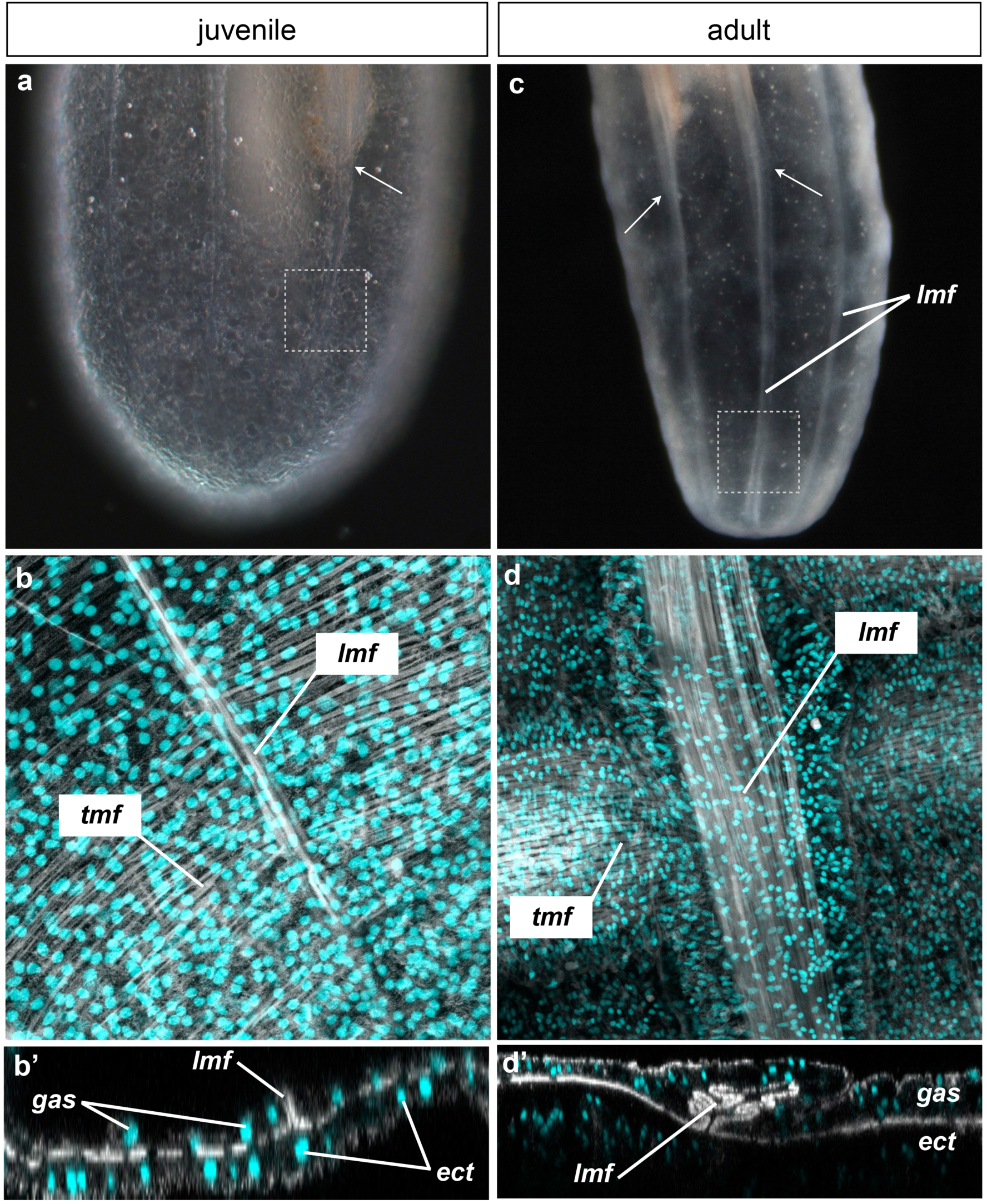
Comparison of the physa anatomy between juvenile (a, b, b’) and adult (c, d, d’). (a) and (c) are macrophotographs. (b, b’, d, d’) are confocal stack images in which the DNA (nucleus) is labeled with DAPI (cyan), and the actin microfilaments (muscle fibers and cell membranes) are labeled with PhallAcidin (white). These confocal stack images are close-ups of the physa where longitudinal muscle fibers are present. (b’) and (d’) are orthogonal confocal projection of the stacks (b) and (d), respectively. White arrows indicate either the end (a) or the continuity (c) of the mesenteries. Dashed squares in (a) and (c) indicate the region represented in (b) and (d) respectively. *ect*, ectoderme; *gas*, gastrodermis; *lmf*, longitudinal muscle fibers; *tmf*, transversal muscle fibers.

### Mesenterial tissue is required for cell proliferation and regeneration

While in the juvenile [physa] no or very little mitotic activity was detected in the majority of cases (Figure 3Ba, a’), 24 out of 63 juvenile [physa] displayed localized cell proliferation at the amputation site (Figure 3Bb-c,b’-d’). Interestingly, DIC and fluorescence imaging revealed that the observed localization and intensity of cellular proliferation were linked to the presence and the amount of remaining mesenterial tissues in the juvenile [physa] (Figure 3Ba’-d’). This observation strongly supports the idea that the presence and amount of mesenteries remaining in the isolated body part, are associated to cell proliferation during regeneration.

In order to directly test our hypothesis that mesenteries are crucial for cell proliferation and regeneration in *Nematostella*, we performed further tissue isolation experiments in adults. By cutting twice transversally, we isolated a part of the mid-trunk region that we named [BE + mes] (Body wall Epithelia *plus* mesenteries). We then opened it longitudinally to separate the body wall epithelia [BE] from the mesenteries [mes], and cultured the [BE] and [mes] separately (Figure 5, Additional file 4: Figure S4). We then analyzed daily if the isolated tissues [BE] or [mes] or the combination of both, [BE + mes], are able to regenerate a fully functional polyp. The [mes] did not regenerate (90 out of 90 cases) but instead, began to fragment one day after isolation, and progressively degraded over time (Figure 5a-c). Surprisingly, while the wound healed, none of the [BE] regenerated neither and they remained intact even after 25 days post isolation (33 out of 34) (Figure 5d-f). Only the [BE + mes] regenerated 10 days after tissue isolation (27 out of 39, Figure 5g-i) and the resulting polyps were able to feed (Additional file 5: Figure S5). To further analyze the importance of the mesenteries to induce regeneration in the surrounding [BE], we performed rescue experiments by grafting [mes] to the endodermal component of the [BE]. Interestingly, nearly half of the grafts regenerated a polyp (6 out of 14 cases, Figure 5j-l). However, regeneration was not complete and none of the regenerated polyps were able to feed. After several weeks, one of them lost its tentacles and all reduced in size (data not shown).

**Figure 5.**
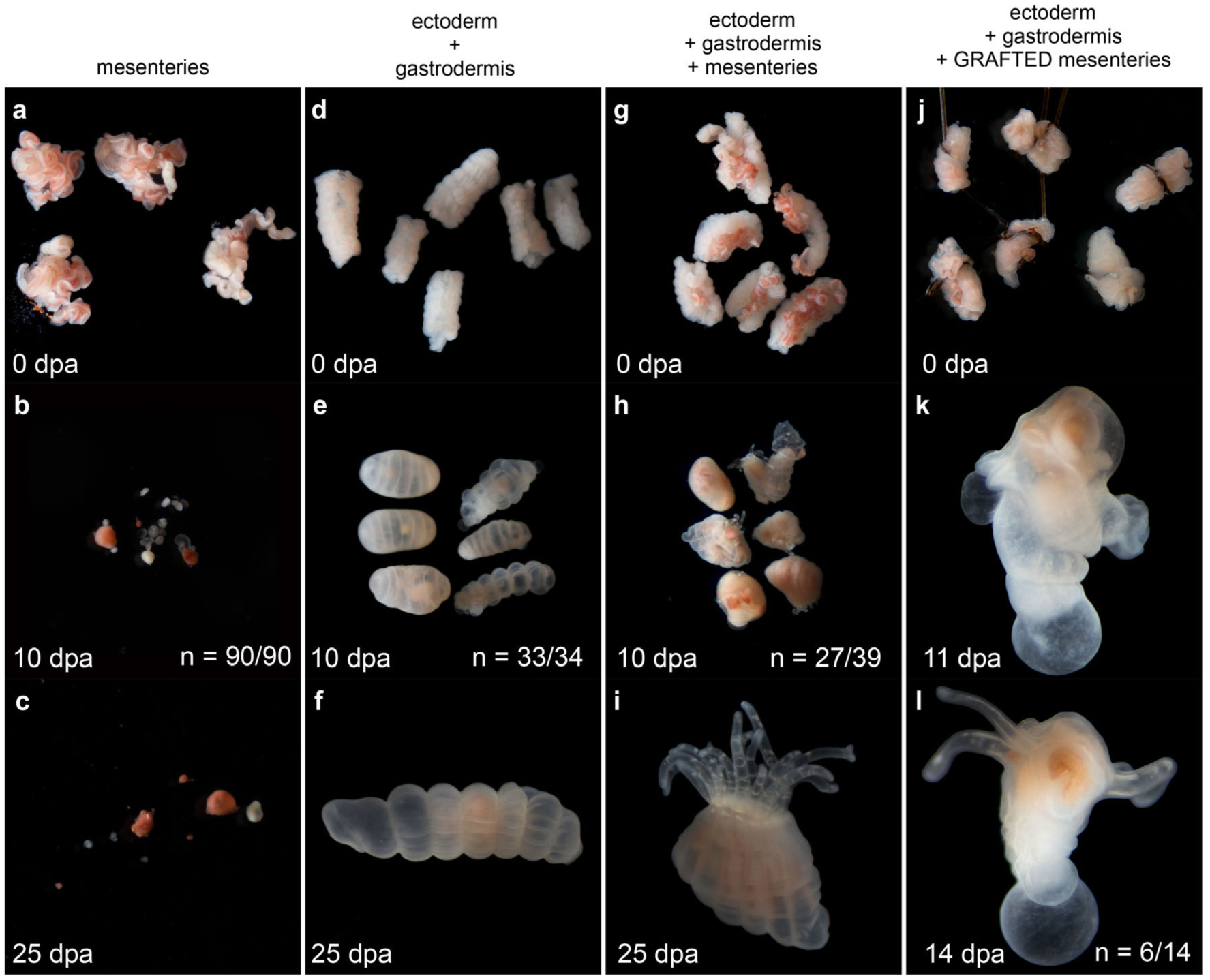
Mesenteries are required for regeneration. Regenerative success was assayed after tissue isolation experiments in which mesenteries were isolated from the ectodermal and gastrodermal epithelia. The isolated tissues or the type of grafting experiment is indicated on top of the panel. The time scale is from top to bottom of the figure and indicated in days post amputation (dpa) in the left bottom corner of each image. White numbers in (b, e, h, k, n) are the n=[number of specimen with represented phenotype]/[total number of analyzed specimen].

To advance our understanding of the role of the mesenteries in inducing regeneration, we analyzed cell proliferation at 24, 72 and 168hpa (7dpa) in the isolated tissues described above. While at 24hpa, we did not observe cell proliferation neither in [BE] nor in [BE + mes], at 72 and 168hpa we detected massive and localized cell proliferation only in the [BE + mes] (Figure 6a-f). At those time points, cell proliferation was detected in a density gradient of EdU+ cells in the tissue, allowing us to determine the site where the future tentacles will form (Figure 6e-f). Taken together, our data clearly show that i) the presence of [BE] is required to maintain the integrity of the mesenteries, and that importantly, ii) the mesenteries are required and possess the capacity to induce cell proliferation as well as regeneration of the body wall epithelia. Thus, a crosstalk between [mes] and [BE] is required for the regeneration process in *Nematostella,* with a particularly crucial role of the mesenteries in the induction of this phenomenon.

**Figure 6.**
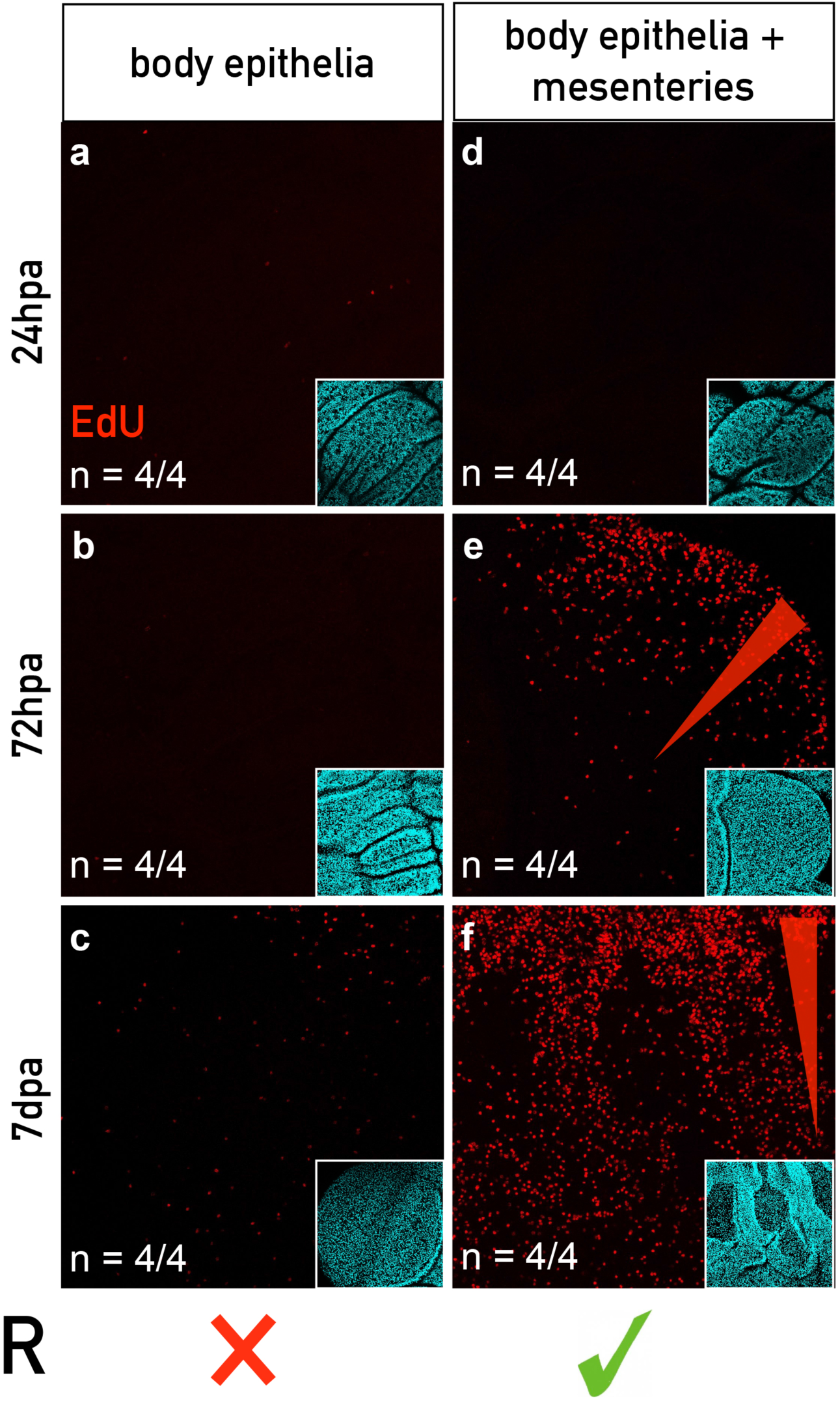
Mesenteries are required for cell proliferation during regeneration. Cell proliferation using EdU staining at various time point during regeneration in adult [body epithelia] (a-c, a’-c’) and [body epithelia + mesenteries] (d-f, d’-f’). Confocal microscopy images showing cellular proliferation (a-f; red) and DAPI (a’-f’; cyan) for corresponding images. Red triangle in (e, f) indicates the gradient of proliferating cells at the regenerating site. No regeneration occurs in the [body epithelia] (a-c, a’-c’) indicated by a red cross at the bottom of the panel in front of R (Regeneration). Regeneration occurs in the [body epithelia + mesenteries] (d-f, d’-f’) indicated by a green check at the bottom of the panel in front of R (Regeneration). n=[number of specimen with represented phenotype]/[total number of analyzed specimen].

### Label retaining cells are present in *Nematostella* tissues

In whole body regeneration models, tissues lacking regenerative capacities have been associated to the absence of undifferentiated stem cells in this compartment (e.g. planarian pharynx [9–11], *Hydra* tentacles [12, 13]). In the previous set of experiments we have identified the juvenile physa that, in contrast to its adult counterpart is devoid of mesenteries, as a body part lacking regenerative capacity. Thus an appealing hypothesis is that the mesenteries are one compartment that acts a as a potential source of adult stem cells involved in regeneration in *Nematostella*.

Quiescence or slow cycling is among the characteristics of vertebrate adult stem cells that participates in the protection of their genomic integrity [64]. In order to identify the existence of slow cycling / quiescent cells populations in *Nematostella*, we carried out label retaining experiments used in many vertebrate models to identify tissue specific stem cell populations [65–69]. After a one-week EdU pulse and extensive rinsing steps in uncut juveniles and adults, the labeled animals were feed once (juveniles) or twice (adults) per week during the 11 weeks (juveniles) or 39 days (adults) periods of the chase (Figure 7).

**Figure 7.**
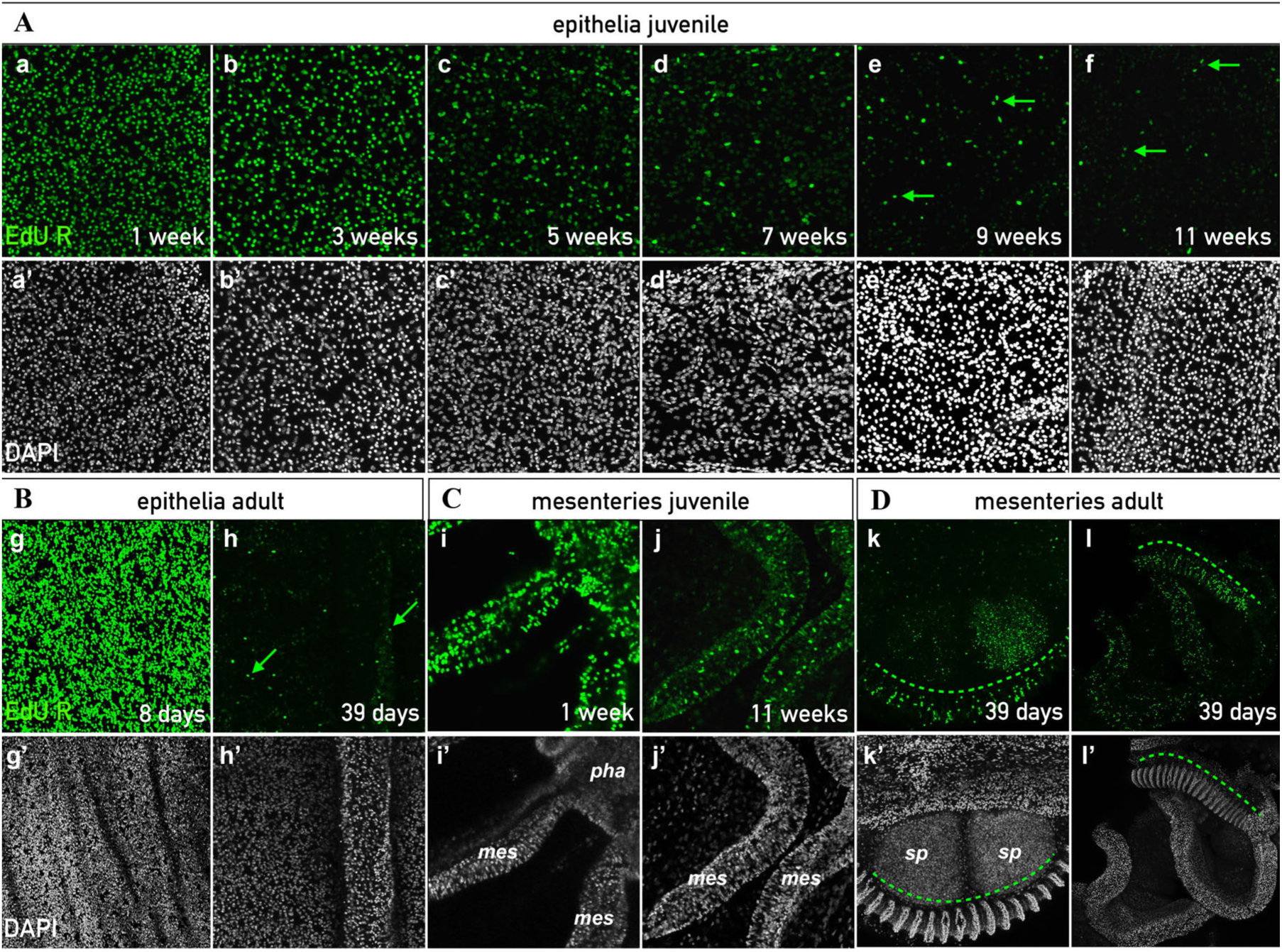
Presence of Label Retaining cells (LRCs) in uncut juveniles (A, D) and adults (B, C). **A**. LRCs present in juvenile epithelia that following a 1 week pulse, were chased at 1 (a,a’), 3 (b,b’), 5 (c,c’), 7 (d,d’), 9 (e,e’), and 11 weeks (f,f’). **B.** LRCs in adult epithelia chased at 7 (g,g’) and 39 days (h,h’) after a 1 week pulse. **C.** LRCs in juvenile mesenteries chased at 1 (i,i’) and 11 weeks (j,j’) after a 1 week pulse. **D.** LRCs in adult mesenteries chased at 39 days (k,k’,l,l’) after a 1 week pulse. LRCs are stained in green (a-l). Nuclei in white (DAPI) (a’-l’). Green arrows (d-h) and the discontinued line (k,k’,l,l’) indicate LRCs pairs and the zone of LRC accumulation in the gonad, respectively. *mes*, mesenteries; *pha*, pharynx; *sp*, sperm mass.

Starting at seven weeks or 39 days after the EdU pulse in juveniles or adults respectively, we began to distinguish isolated and randomly localized EdU+ label retaining cells (LRCs, Figure 7). Those LRCs were detected as expected in the mesenteries of juvenile and adult polyps (Figure 7C,D). Note that in addition to randomly dispersed LRCs in adult mesenteries, we also observed dense EdU+ cells in a restricted region close to the gametes, suggesting the presence of germ line precursors in this specific location (Figure 7D, Additional file 6: Figure S6g,g’).

To our surprise we also detected LRCs within the body wall epithelia (Figure 7A,B. Interestingly, we detected LRC pairs in juveniles and adult epithelia, suggesting that a subpopulation of quiescent cells are able to divide to maintain tissue homeostasis in *Nematostella* (Figure 7Ae,e’,f,f’,Bh,h’). In adults, the density of LCRs after 39 weeks seemed reduced in the endoderm compared to the ectoderm, suggesting that the cellular renewal takes place faster in the endoderm than in the ectodermal (Additional file 7: Figure S7a,a’,b,b’). To reinforce the idea that we have detected quiescent/slow cycling cells in *Nematostella*, LRCs were still detected in the epithelia of the body wall (as well as in the mesenteries) five months after the initial EdU pulse (Additional file 8: Figure S8k,k’). However, while after 39 days of chase LRCs were still detected in the tentacles, after 5 months no more LRCs were observed in this part of the body (Additional file 8: Figure S8h-j,h’-j’), suggesting that the cellular turnover (causing the elimination of the LRCs) in the tentacle region is increased compared to the body column.

To make sure that the LRCs we detected are not all terminally differentiated cells, we analyzed the nuclear morphology and organization of those EdU+ cells. While we detected differentiated cells such as cnidocytes, excretory cells and batteries of nematocysts (Additional file 7: Figure S7d-f,d’-f’), we also observed a subpopulation of cells with highly condensed DNA (Additional file 7: Figure S7c,c’), reminiscent of stem cell nuclei in *Hydractinia* [22]. The presence of LRCs in the mesenteries (other than potential germ line precursors) as well as in the epithelia that are able to re-enter the mitotic cycle under physiological conditions, further support the presence of adult stem cells in the anthozoan *Nematostella*.

### Irradiation resistant LRCs in the mesenteries and response to the amputation stress

The capacity to retain EdU labelling is one feature of slow cycling/quiescent stem cell populations that thus represent an increased resistance to irradiation. Another characteristic is the ability to re-enter the cell cycle after a challenge (e.g. amputation stress) [64]. In order to test this ability in *Nematostella*, we inhibited mitosis in uncut animals using X-ray irradiation. As expected, no more cell proliferation was detected four hours after irradiating the animals at 100Gy (Additional file 9: Figure S9a,a’,b,b’). In order to analyze if mitotic activity can re-emerge in the irradiated animals, we performed EdU incorporation at various time points following irradiation on uncut polyps (Additional file 9: Figure S9c-e,c’-e’). Interestingly, we observed EdU+ cells starting at 96hours post irradiation (hpirr) that were dispersed randomly within the mesenteries and the basal part of the pharynx but not at all in the epithelia (Additional file 9: Figure S9e,e’). These data clearly show that a pool of cells within the mesenteries and the basal part of the pharynx is able to escape the effects of irradiation.

In order to test if this small pool of irradiation resistant quiescent/slow cycling cells is sufficient to allow proper regeneration, we performed a sub-pharyngeal amputation four hours after irradiation. Doing so, we observed that regeneration was impaired. More precisely, the majority of irradiated animals were blocked prior to the formation of the pharynx at step 2 of the sub-pharyngeal oral *Nematostella* regeneration staging system [62] (100Gy: Figure 8A, Additional file 10: Figure S10i,i’; 300Gy: Additional file 11: Figure S11b,b’, Additional file 12: Figure S12). To analyze the capacity of the quiescent/slow cycling cells to respond to the amputation stress, we repeated the same experiment (sub-pharyngeal amputation of irradiated polyps) that was followed by EdU labeling at various time points post dissection. To our surprise, we detected the presence of EdU+ cells in the most oral part of the mesenteries already at 48hpa in the amputated polyps (Figure 8Be,e’). Later (120hpa), in addition to the ones in the oral tips of the mesenteries, we also observed EdU+ cells in the epithelia at the amputation site (100Gy: Additional file 10: Figure S10i,i’; 300Gy: Additional file 11: Figure S11b,b’). While quiescent/slow cycling cells that escape irradiation are able to re-enter the cell cycle in uncut as well as cut polyps, their timing of emergence and dispersion is different. They are either detected 96hpa and distributed along the oral-aboral axis of the mesenteries in uncut animals, or restricted to the oral most part of the mesenteries in dissected polyps starting 48hpa. Latter observation suggest that quiescent/slow cycling cells are migrating from the aboral region of the mesenteries towards the oral most part of those structures in response to the amputation stress.

**Figure 8.**
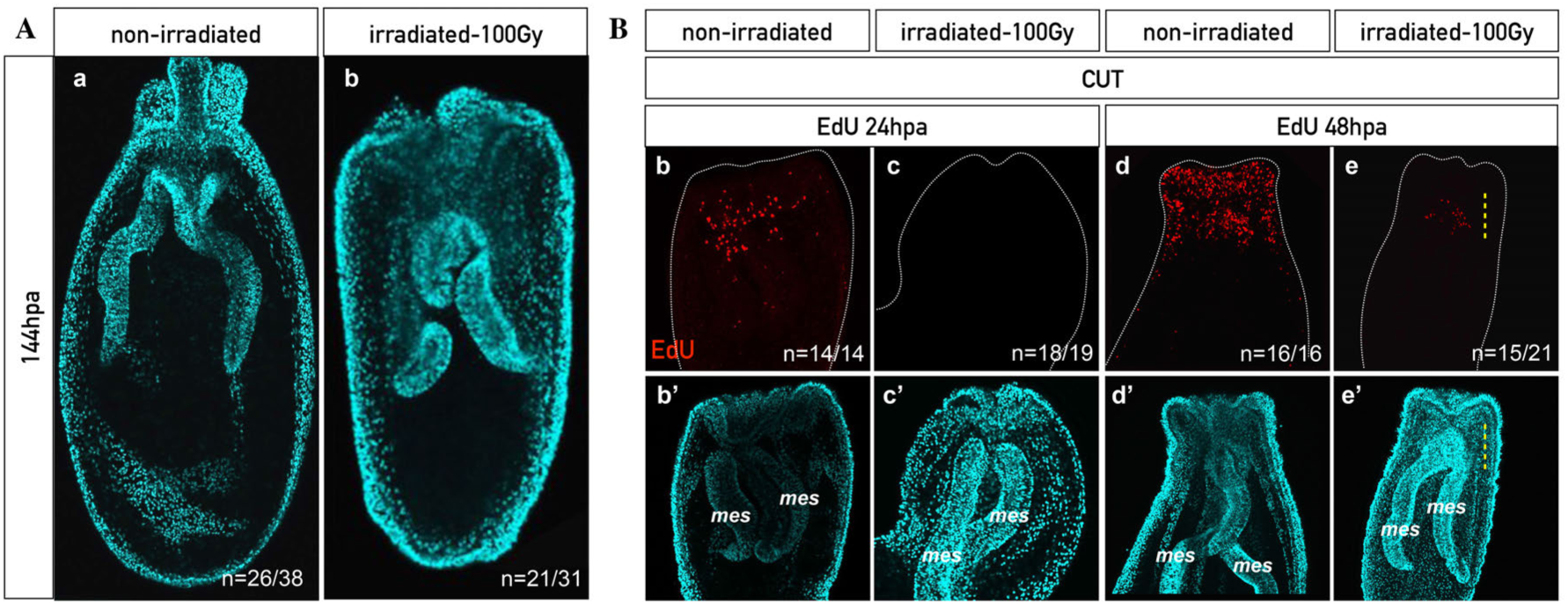
**A.** X-ray irradiation perturbs head regeneration. Confocal images of control non-irradiated (a) and X-ray irradiated (a’) animal at 144hpa. Nuclei are in cyan (DAPI) (a, a’). **B.** A population of cells in the mesenteries is able to re-enter the mitotic cycle 48 hours post irradiation (hpirr). Confocal images of control non irradiated (b,b’: 24hpa; d,d’: 48hpa) and irradiated (c,c’: 24hpa; e,e’: 48hpa) polyps. A few cells in the mesenteries, localized at the amputation site, were able to re-enter the cell cycle between 24 and 48hpa (e, e’). EdU staining in red shows cell proliferation (b-e). Nuclei are in cyan (DAPI) (b’-e’). *mes*, mesenteries; *pha*, pharynx; *ten*, tentacles. n=[number of specimen with represented phenotype]/[total number of analyzed specimen].

Taken together these results show that i) the quiescent/slow cycling cells of the mesenteries alone are not sufficient to allow full regeneration, ii) that at least an additional pool of fast cycling cells (the ones affected by irradiation) are required for reformation of lost body parts and iii) that the LRCs of the mesenteries that escape the effects of irradiation, are activated by the amputation stress by displaying mitotic activity and by migrating towards the amputation site. Thus, we now provide a series of evidences that highlight the presence of quiescent/slow cycling stem cells in *Nematostella*, in particular, those located within the mesenteries.

### LRCs are migrating from the mesenteries to the body wall epithelia

We have shown that a crosstalk between the mesenteries and the epithelia is required for cell proliferation/regeneration and that quiescent/slow cycling stem cells are potentially migrating from the mesenteries to the surrounding epithelia in response to the amputation stress. To test this hypothesis, we designed a protocol that combined EdU pulse and chase with grafting experiment. More precisely, we labeled LRCs of the entire polyp and isolated its mesenteries, named [mesLRC] (LRC stained mesenteries). We then grafted the [mesLRC] to the endodermal component of the [BE] from an unlabeled animal (Figure 9A). After 14 days, 3 out of 18 grafts regenerated. Nonetheless, all 18 grafts were fixed in order to chase the EdU staining and determine the localization of LRCs in the chimeric animals. To our surprise, we not only observed LRCs in the previously unlabeled [BE] (5 out of 18 cases), but also far from the [mesLRC] source (Figure 9Ba). In one of the regenerated [mesLRC] [BE] grafts, a large amount of LRCs have even migrated towards the newly formed tentacles (Figure 9Bd). This clearly demonstrates that LRCs are able to migrate from the mesenteries towards the epithelia, and even within the epithelia to integrate specific regions. Importantly, we also observed LRC pairs (in addition to single LRCs), showing that the LRCs that migrated from the [mesLRC] to the [BE] were able to divide and are not post-mitotic (Figure 9Bb,c). A detailed analysis of the DNA of migrating LRCs using confocal microscopy revealed that i) their nucleus is round, ii) very similar in size, iii) posses a compact DNA and iv) the majority is located in the endoderm of the [BE] (Figure 9C). Latter observation strongly suggests that the graft re-establishes the physical contact between the endodermal components of the mesenteries and the [BE]. In sum, we now have clear evidence that LRCs, or their immediate progeny, are able to migrate from the mesenteries towards the body wall and tentacle epithelia under stress conditions where they may participate in regenerating missing body parts.

**Figure 9.**
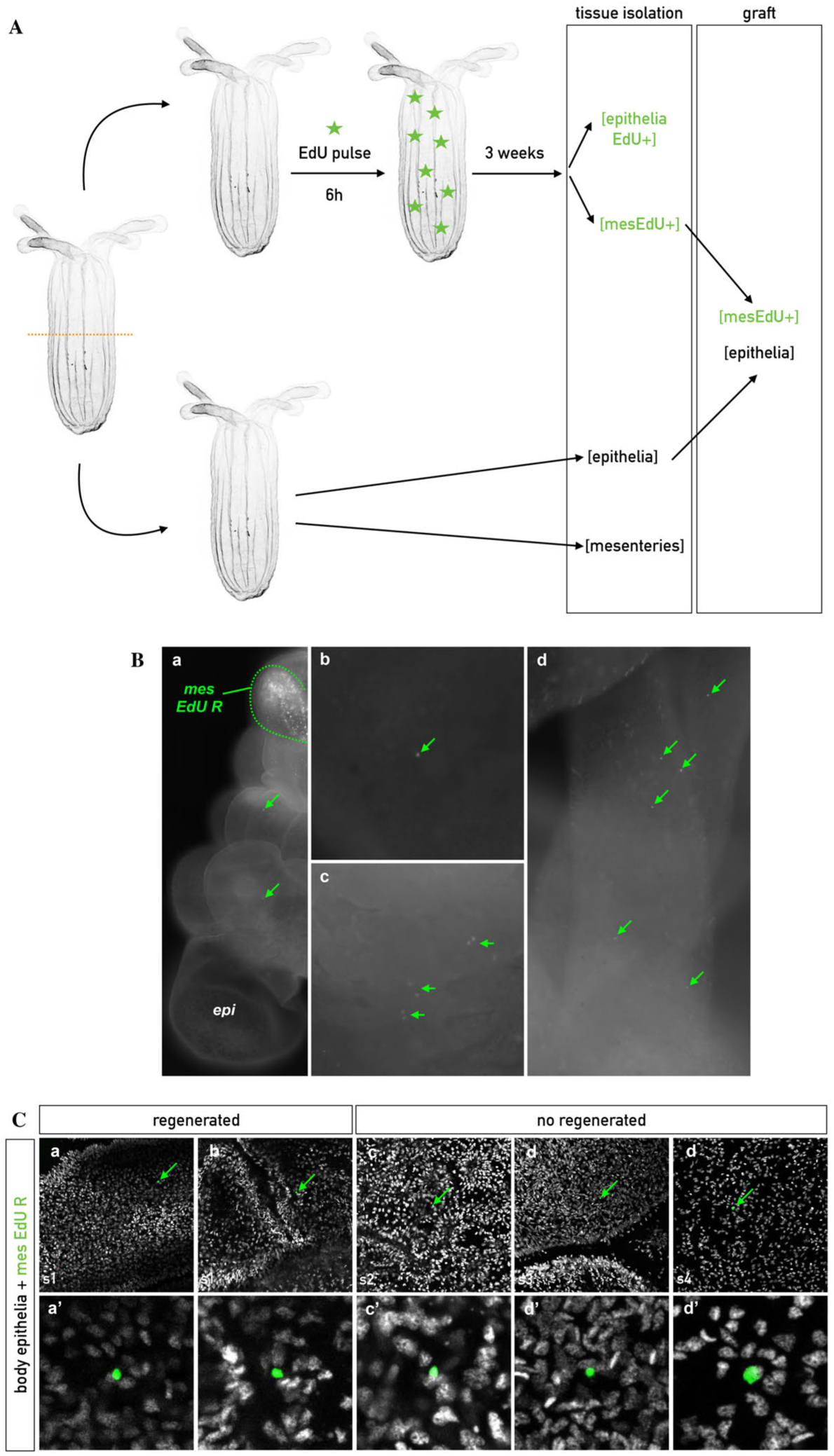
**A.** Diagram of the protocol to assess cell migration in graft experiments. [epithelia EdU+] and [mes EdU+] represent the isolated epithelia of the body wall and the mesenteries, respectively, that were labeled with EdU (from clone 1). [epithelia] and [mesenteries] represent the isolated epithelia of the body wall and the mesenteries, respectively, that were not labeled (from clone 2). **B.** Fluorescence imaging of the resulting graft experiments ([epithelia] + [mesLRC]). The discontinued green line indicates the position of the [mesLRC]) 14 days after being grafted into the [epithelia] (*epi*). Green arrows indicate EdU+ cells from the [mesLRC] that retained the EdU and migrated into the non labeled [epithelia] (a and b). In addition, they were able to divide as indicated by the presence of cell pairs (c) that can be localized in the regenerating tentacles (d). **C.** Confocal imaging of EdU+ cells from the [mesLRC] that retained EdU and migrated into the non labeled [epithelia]. Overlap of the nucleus staining in white (DAPI) and EdU R in green (a-d, a’-d’). a’-d’ are magnification of the EdU+ cell from a-d.

### LRCs localized in the epithelia respond to the amputation stress

In addition to the clearly defined pool of mesenterial LRCs, we have also observed LRCs within the epithelia that were however, less numerous compared to the mesenterial ones (Figure 7). To perform a detailed characterization of all LRCs present in *Nematostella*, we also analyzed the dynamics of LRCs localized in the epithelia of the body wall during regeneration. We performed sub-pharyngeal amputation on LRCs positive adult polyps (Figure 10A) and analyzed at various regeneration time points (24-192hpa) i) the localization of the LRCs, ii) their ability to divide and form clusters in response to the amputation stress and iii) their becoming in the newly regenerated head (Figure 10B). The LRCs of the mid-body and aboral regions remained largely unaffected. However, we detected a general, although variable, accumulation of LRCs as well as the presence of LRC clusters at the amputation site starting at 24hpa (Figure 10Ba-d,a’-d’). This variability could be either associated i) to the different metabolic states of each individual or ii) to the increased division rate at the amputation site at later stages that may cause the dilution of the EdU signal or iii) both. Importantly though, we detected single LRCs as well as pairs of LRCs in the newly formed tentacles 8 days post-amputation (192hpa) (Figure 10Be-e’’).

**Figure 10.**
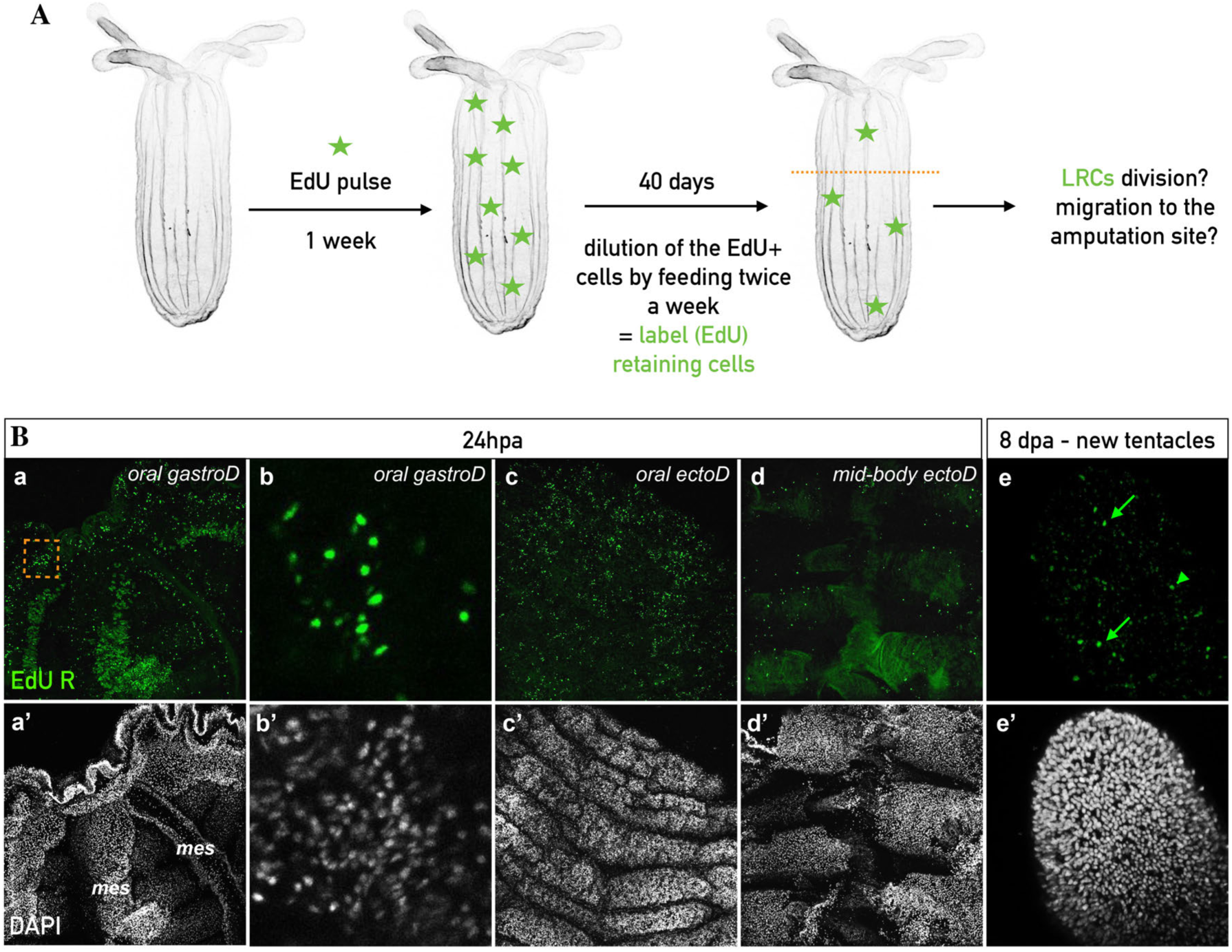
**A.** Diagram of the protocol assessing LRC dynamics during regeneration. After a one-week pulse with EdU, adult polyps were rinsed thoroughly with 1/3x seawater and kept for 40 days. Then, sub-pharyngeal amputation was performed and LRCs were chased at 24, 48 and 72hpa. **B**. Confocal imaging of LRC dynamics in regenerating polyps. LRCs are stained in green (a-e). Nuclei are in white (DAPI) (a’-e’). Regenerating adult polyps at 24hpa (a-d, a’-d’), accumulation of the LRCs cells is detected at the oral side (a,b,d) but not in the epithelia of the body wall (c). b and b’ are the magnification of the square in (a) that indicate a cluster of LRCs at the amputation site during the regeneration process. LRCs are detected in the newly regenerated tentacles (e,e’; green arrows). *mes*, mesenteries; *mid-body ectoD*, mid-body region ectodermal view*; oral ectoD*, oral region ectodermal view*; oral gastroD*, oral region gastrodermal view.

Taken together, our data show that the LRCs of the epithelia, although they are not able to escape the effects of irradiation, i) are activated in response to the amputation stress by dividing and accumulating at the amputation site and ii) participate in the newly formed head (e.g. tentacles epithelia), providing clear evidence that those epithelial LRCs are actively involved in tissue renewal and thus, also possess stem cell-like characteristics.

### Accumulation of fast cycling cell at the amputation site during oral regeneration

We have seen that the mesenterial LRCs (reactivated during regeneration following irradiation) alone are not sufficient to promote full regeneration. This shows the necessity of a tissue crosstalk and in particular, the requirement of another pool of (stem) cells localized in the body wall epithelia. During the irradiation experiment, we blocked cell proliferation in cells that are undergoing mitosis including fast cycling cells required for tissue homeostasis. In order to test if a population of fast dividing cells from the uncut animal also migrates to the wound site during regeneration, we performed a 1h EdU pulse in uncut animals followed by extensive washes to eliminate any excessive EdU. This EdU pulse labels the fast dividing cells involved in tissue homeostasis (Figure 11Aa). We then performed a sub-pharyngeal amputation and chased the EdU+ cells during the process of regeneration from 0 to 120hpa (Figure 11A,B).

**Figure 11.**
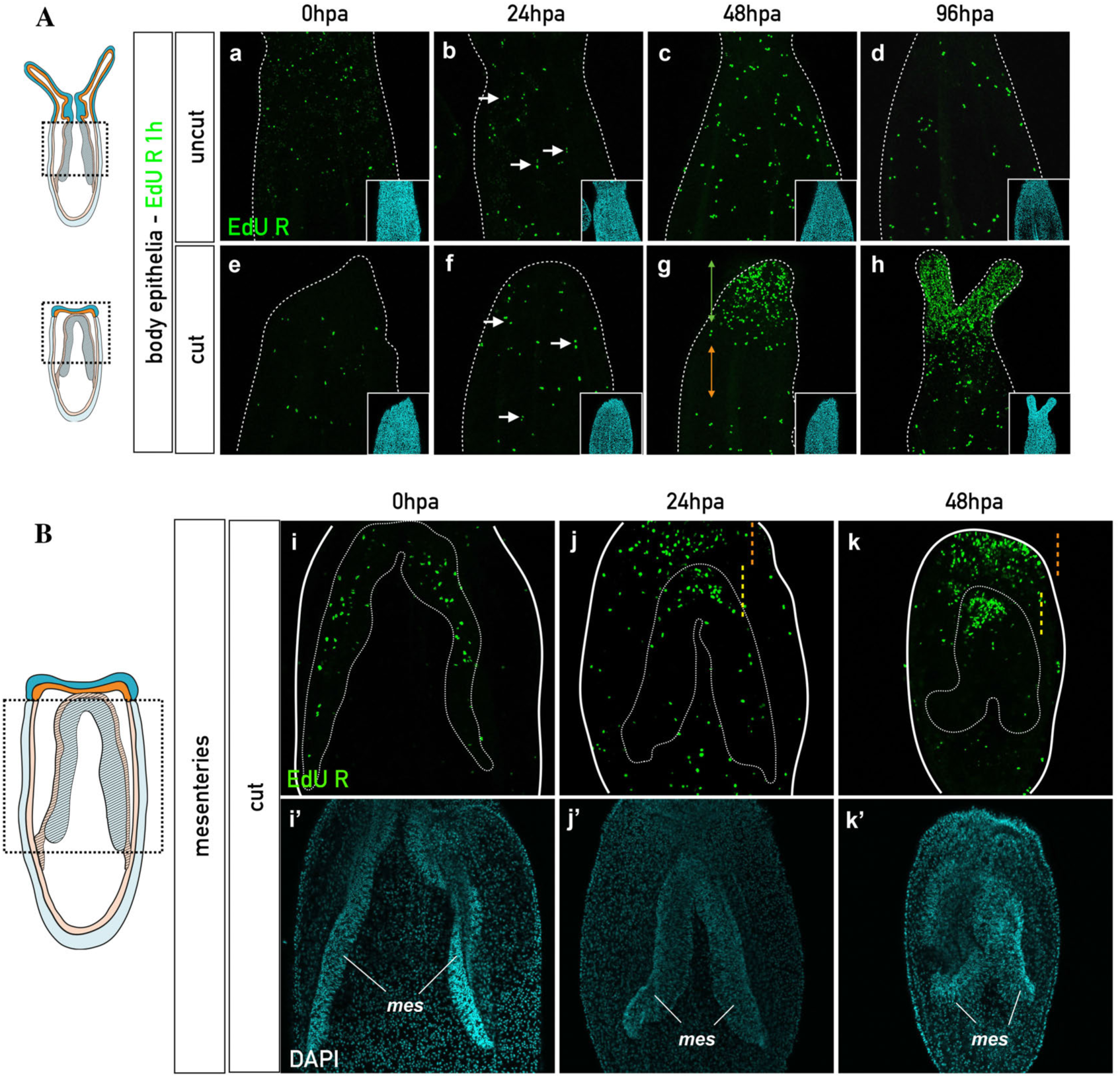
Accumulation of EdU+ cells at the amputation site during regeneration. Confocal images of the EdU “pulse-1h-and chase experiment” on uncut (epithelia: Aa-d) and cut (Epithelia: Ae-h; Mesenteries: Bi-k) polyps analyzed at 0, 24, 48 and 96 hours post-pulse (hpp). In A and B, nuclei are labeled with DAPI (cyan) and EdU retaining cells are recognized by their green staining. White arrows at 24hpp in uncut (Ab) and cut (Af) polyps indicate the appearance of EdU cell pairs. Green and orange double arrowheads in Ag indicate the accumulation of the EdU+ retaining cells at the amputation site of the regenerating polyp, and the EdU depleted zone, respectively. No accumulation of EdU+ retaining cells is detected in uncut animals (Aa-d). In B, the accumulation of EdU+ retaining cells in the most oral part of the mesenteries starts at 24hpa (Bj) and become stronger at 48hpa (Bk). The green double arrowhead and the discontinued line, in Bj and k, indicate the accumulation of the EdU+ retaining cells, in the epithelia and in the mesenteries, respectively. The orange double arrowhead in Bk indicates the EdU depleted zone in the mesenteries. *mes*, mesenteries.

In uncut control animals, EdU+ cells were randomly dispersed throughout the body wall epithelia as well as in the mesenteries for all analyzed time points [59](Figure 11Aa-d, Additional file 13: Figure S13). Starting at 24 hours post-pulse, first homeostatic EdU+ cell divisions are detected (still in a dispersed manner) in uncut animals, as indicated by the presence of EdU+ cell pairs (Figure 11Ab-d). Interestingly, in sub-pharyngeally amputated polyps, we observed a strong accumulation of EdU+ cells in the oral regions of the mesenteries while their remaining aboral parts progressively become devoid of those cells (Figure 11B). This accumulation, indicating amputation-site-directed cellular migration within the mesenteries is already visible at 24hpa Figure 11Bi,j, while EdU+ cells are still randomly located within the epithelia (Figure 11Ae,f). However, starting at 48hpa, EdU+ cells accumulate massively in a restricted region of the epithelia at the amputation site and, in a few cases (48hpa: 9 out of 31), a zone depleted of EdU+ cells is observed in the region below the massive accumulation of EdU+ cells (Figure 11Ag,h).

Those observations are further quantitatively supported by carefully counting individual Edu + cells in three different zones (Z1’, Z1’’ and Z2) of the body epithelia along the oral-aboral axis of the uncut *vs* cut animal during regeneration (Additional file 14: Figure S14). In addition to the difference in the average number of EdU+ cells in zone Z1’ in uncut (sub-pharyngeal region) *vs* cut (amputation site), this cell counting revealed clear differences in the Z1” (lower mid-body region) and Z2 (aboral end) zones for the same conditions. In fact, while the average number of EdU+ cells in Z1” and Z2 is constant or decreasing respectively in regenerating animals, it is increasing in both zones in uncut controls (Additional file 14: Figure S14). In sum, these results show that fast cycling cells (required for regeneration as eliminated during irradiation) from the region below zone Z1’ of the body wall epithelia, migrate towards the amputation site and in cooperation with LRCs, actively participate in the renewal of missing tissues.

## Discussion

In the present study, we have highlighted the unexpected role of the mesenteries not only as a reservoir of quiescent/slow cycling stem cells, but also as the tissue harboring the signal to induce cell proliferation and regeneration in *Nematostella*. Long-term EdU pulse and chase experiments revealed the presence of quiescent/slow cycling cells (LRCs) that respond to the amputation stress by reentering the mitotic cycle, accumulation at the wound-site and participate in the reformation of missing structures. Combining classical graft experiments with EdU pulse and chase experiments, revealed cellular migration from the mesenteries to the epithelia, as well as toward the amputation site in response to the amputation stress. We further showed that LRCs alone are not sufficient and that homeostatically fast cycling cells are required for regeneration. In sum, our present work has not only revealed the requirement of a tissue crosstalk between the mesenteries and the epithelia of the body wall to initiate the cellular dynamics underlying regeneration but importantly also the synergic effect of fast cycling cells and slow/quiescent stem cells to enable full reformation of lost body parts. This study provides also for the first time in an anthozoan cnidarian a set of strong evidences for the presence of adult stem cell populations that are directly involved in the regenerative response.

We thus propose a mechanistic model for head regeneration in *Nematostella,* in which the mesenteries, once in contact with the epithelia of the amputation site (Step 1, [62]) emit a regeneration-initiating signal that activates fast and slow-cycling stem cell populations from the mesenteries as well as the epithelia by inducing their division and migration toward the wound site. Importantly, a synergic effect between those two stem cell populations is required to pursue and complete the regeneration process. The regeneration inducing signal emitted by the mesenteries, the mechanism of synergic cooperation between the two stem cell populations and whether de- or trans-differentiation processes are also involved in the tissue repair and regeneration program of *Nematostella*, remains to be investigated.

### Distinct cellular mechanisms between oral vs aboral regeneration in Cnidaria

By comparing the regenerative capacity of various body parts to reform missing oral as well as aboral structures, we observed that regeneration of aboral tissues does not involve massive cellular proliferation at the amputation site, while oral regeneration does. These data suggest that aboral regeneration either i) uses a different cellular mechanism than the one described for oral regeneration in order to reform the missing physa and/or ii) that the requirement of regenerating aboral tissues is less urgent (and thus does not require massive localized cell proliferation) than quickly reforming a missing head region that is more complex and crucial for feeding and defense. A distinct cellular mechanisms for oral *vs* aboral regeneration has been recently described in the cnidarian hydrozoan *Hydra* and *Hydractinia* [20, 22] supporting the idea that using different mechanism to regenerate opposite part of the body in a same system seems to be a conserved feature among Cnidaria. The studies performed in Hydrozoa show that similar to our observations, no massive cell proliferation is detected at the aboral amputation site [20, 22]. In *Nematostella*, while we also observe a slight reduction in tentacle size during aboral regeneration that could contribute to the reformation of the physa via tissue reorganization, we have no evidence for a complete transformation of the body column into an aboral fate prior to re-growing a fully functional polyp. However, recent work on *Nematostella* shows a clear difference in the transcriptional response of the oral *versus* aboral regeneration process [61]. This study also shows that the early molecular response (8hpa) to the amputation stress is more similar between oral *vs* aboral regions, compared to later time points (24 and 72hpa) [61]. Interestingly, our previous study on *Nematostella* regeneration revealed that early oral regeneration steps (from step 0 to step 1; between 0 to 24hpa) are cell proliferation independent, while the later ones (from step 2 to step 4, between 24 to 120hpa) are cell proliferation dependent [62]. Thus, in *Nematostella*, the mechanisms driving the early steps of tissue regeneration such as wound healing seem similar between oral *versus* aboral regeneration as they are both cell proliferation independent. However, additional experiments are required to gain a better understanding of the cellular and molecular mechanisms involved in aboral regeneration in *Nematostalla,* in particular during later regeneration phases, when initial wound healing and early response is completed.

### Regenerative capacity of a given body part is age-dependent

It has been previously shown that oral regeneration after sub-pharyngeal amputation in *Nematostella,* occurs at a comparable time-scale in both juveniles and adults and that cellular proliferation is required in both cases [62]. Our present study revealed that clear variation in the capacity to regenerate exists in one given body part depending on the age of the organism. The obtained data show that the tissue composition within a given body region is changing with age. In particular, the isolated physa regenerates only in adults, but not in juveniles that are unable to induce sufficient cellular proliferation, probably by the lack of i) stem cell population(s), ii) initiation signal(s) or both. In addition, this approach was useful to identify which body part/structure of *Nematostella* is required for the initiation of regeneration and to orient our research on the potential stem cell population(s) involved in the regeneration process.

### Mesenteries are required and sufficient to induce cell proliferation and regeneration

Taking advantage of the differential regenerative capacity of juvenile and adult physa, we identified the mesenteries as crucial structures to induce cellular proliferation and subsequent regeneration in the surrounding tissues. By performing tissue dissociation and grafting experiments, we have shown that isolated mesenteries, grafted back to the remaining [BE] of the same individual, are sufficient to induce regeneration. Extending this finding to other body parts of juveniles that did not regenerate when isolated such as [tentacles], [tentacles + half pharynx], [tentacles + full pharynx] and [pharynx], it seems plausible that the common factor explaining the lack of regeneration in those juvenile body parts are the missing or not completely formed mesenteries. [tentacles] or [tentacles + half pharynx] isolated from adults did not regenerate neither, while [tentacles + full pharynx] or [pharynx] did. Tentacles clearly lack mesenterial tissues and interestingly, the mesenteries are anchored to the pharynx in its aboral regions. As the half pharynx that remained with the tentacles in [tentacles + half pharynx] corresponds to the oral half of the pharynx, the absence of regenerative capacity in juvenile and adult [tentacles] or [tentacles + half pharynx] can thus be linked to the absence of mesenteries in those body parts. Fully formed mesentery anchors are present in the adult pharynx that are sufficient to induce regeneration from [tentacles + full pharynx] or [pharynx]. In analogy to our findings in the juvenile physa, we expect that the mesenterial anchors, or at least important components of it (i.e. stem cells), in the juvenile pharynx are not fully formed or are absent. This may explain the absence of regenerative capacity of those isolated body parts.

One puzzling observation in the graft experiments (restoring the regenerative capacity) was that the regenerated animals possessed a tentacle crown but not a mouth and were therefore unable to feed. One probable explanation for this is the number of mesenteries required for full regeneration. In the graft experiment performed in our present study only one mesentery has been grafted into the epithelia leading to a partial regeneration (head without mouth/pharynx). By testing the removal of only 7, 6 or 5 mesenteries on 8 total from the adult body wall epithelia, we have observed that at least two mesenteries are required for proper mouth formation. In none of the cases in which only one mesentery was left in contact with the body wall epithelia, the regenerated polyps were able to fed, showing that only partial regeneration occurs (data not shown). This evidence for the requirement of at least two mesenteries for proper pharynx reformation is further enhanced by previous observations that during oral regeneration the two mesenteries fuse to one to each other, and that the fused oral parts of the mesenteries give rise to the vast majority of the newly formed pharynx [62].

### A tissue crosstalk is required for initiating a regenerative response

The observation that the mesenteries are required to induce regeneration in the surrounding epithelia leads to three main hypotheses concerning the mechanism involved: 1) A long-range diffusing “regeneration” signal emitted by injured mesenteries is required to initiate cellular proliferation in the surrounding epithelia. 2) The physical integrity of the tissue that links the mesenteries to the body epithelia is important to relay the “regeneration” signal. 3) A pool of stem cells (or their offspring) migrates from the mesenteries to the body wall epithelia to initiate the regeneration process. We tested the first hypothesis by incubating the isolated body wall epithelia in close but not physical contact with isolated pieces of mesenteries in the same wells. In none of the cases the body wall epithelia regenerated (data not shown), suggesting that the molecular induction signal (if there is any) from the mesenteries is not sufficient alone, or that it is too diluted in the well, to induce regeneration of the epithelia. Alternatively, it might also be possible that only endodermal cells respond to the induction signal to transform it into a regenerative response. In this case, the fact that the isolated epithelia wounded and closed with the endodermal cells inside prevents the signal to target the responsive cells. While additional experiments are required to fully address hypothesis 1, we currently favor hypothesis 2 and 3 to explain the role of the mesenteries in the regeneration process of *Nematostella*.

The importance of physical tissue crosstalk during regeneration has been studied during vertebrate intestinal epithelium regeneration in which the connective tissues have been shown to be crucial for the regeneration of the epithelium [70]. In addition, cell/cell contact molecules such as integrins are known to be involved in the regeneration process in vertebrates [71, 72] [73]. The migration of stem cells to the amputation site has been shown for regeneration of complex structure [74, 75] and it has been proposed that mechanical stress induces cell migration as in smooth muscles [76]. Our study has provided clear evidences for a cellular transfer of cells from the mesenteries to the epithelia as well as the existence of two populations of cells that have stem cells characteristics (Hypothesis 3). The observation that the mesenteries disintegrate when isolated from the body wall epithelia highlight the importance of a tight and physical tissue interaction between the epithelia and the mesenteries to maintain the integrity and/or homeostasis of the mesenteries (Hypothesis 2). Hypothesis 2 and 3 are further reinforced by the fact that the process of regeneration is delayed in graft experiments compared to [BE + mes] (14 *vs* 10 days, respectively). This could be explained by the fact that the tissue connecting the mesentery and the body wall epithelia [62] needs to be first reformed, before either the inductive signal can be relayed, or before the stem cell population can migrate from the mesenteries to the body wall epithelia.

### Migration of different pools of cells during regeneration

Our work highlights evidences that cell migration occurs during *Nematostella* regeneration as it is the case in others invertebrate and vertebrate regeneration models studied so far [1]. These evidence are: i) LRCs migrate from the mesenteries to the surrounding epithelia, ii) fast cycling cells involved in tissue homeostasis migrate to the amputation site, and iii) following irradiation, slow cycling EdU+ cells within the mesenteries that escape irradiation accumulate the most oral part of the mesenteries, while EdU+ cells are randomly dispersed in the mesenteries in irradiated uncut animals. These data show that this population of cells re-entering the cell cycle in irradiated animals migrate to the amputation side during regeneration. However, latter cell population is not sufficient alone to fully complete regeneration as the regeneration process in irradiated polyps is blocked at step 2 of the sub-pharyngeal *Nematostella* staging system [62]. We recently showed that inhibition of cell proliferation using Hydroxy Urea during *Nematostella* regeneration, blocked the regeneration process at step 1 [62], suggesting that the slow cycling, irradiation-resistant cells that re-entered the mitotic cycle in response to the amputation stress are crucial to transition at least from step 1 to step 2.

In *Nematostella*, although we were able to detect a “first wave” of migrating cells only between 24-48hpa, we cannot exclude that cell migration of other cell populations also occurs at earlier time points between 0 to 24hpa, as it has been shown in *Hydra* [20]. The potential second “wave” of cell migration, reflected by the decrease in the number of the EdU+ cells in zone Z2 (most aboral part of the animal) during regeneration, might correspond to a replacement of stem cells populations in a zone that got depleted of it. This kind of stem cell replenishment has been shown during tissue homeostasis or tissue transplantation experiments in *Hydra* to repopulate a tissue that was i-cell deficient [77] [78]. Our experiments show that fast as well as slow cycling cells are able to migrate during *Nematostella* regeneration, however, we currently don’t know if the migrating cells are the stem cells or their progeny. Further characterization of the molecular identity of those migrating cells as well as the development of *in vivo* tools is required in *Nematostella* to answer this question.

### Stem cells in Anthozoa

Prior to the present study, cells populations with clear stem cell characteristics have not been described in *Nematostella* or more generally in anthozoan cnidarians (sea anemones, corals) [32,57,59]. Our present work highlights the existence of two populations of proliferating cells that are synergistically required for regeneration: i) one pool, composed of slow cycling/quiescent cells (LRCs) and ii) another pool, composed of fast cycling cells involved in tissue homeostasis in the uncut animal. Both cell populations possess features that are characteristic for stem cells: 1) The behaviour of the slow cycling/quiescent/label retaining cells (LRCs) *per se*, as it is one important feature of adult stem cells that is deployed in mammalian stem cells to preserve genomic integrity [64, 69]. These slow cycling/quiescent cells are able to escape X-ray irradiations and respond to the amputation stress by migrating and accumulating towards the wound site, where they actively divide and participate in the reformation of lost structures. While the potency of those LRCs has not been properly addressed yet, we already observed that they are able to give rise to sensory cells within the regenerated tentacles. 2) The fast cycling cells, respond to the amputation stress by migrating towards the amputation site where they actively divide and are required for regeneration, probably for later regeneration steps (3 and 4). In the hydrozoans *Hydra* and *Hydractinia*, fast cycling stem-cells (i-cells and/or their progeny) also divide and migrate to the amputation site and actively participate in the regeneration process [20, 22]. Thus, our results suggest a conserved pool of fast cycling stem cells in cnidarians that are involved in tissue homeostasis and regeneration.

Stem cell/pluripotency markers in ctenophores and cnidarians are known to include the germ lines markers *piwi*, *vasa*, *nanos* and *PL10* [22,79–84] [85]. A previous study in *Nematostella* analyzed the expression pattern of those genes (with the exception of *piwi*) from early development to the primary polyp stage [86]. In the juvenile, *Nvnos1* is found in some patches of cells in the ectodermal epithelia of the body wall. The authors suggested that these patches of *Nvnos1*+ cells are “population of nematocyst precursors with stem cells characteristics” [86]. However, all other analyzed germline genes *NvPl10*, *Nvvasa1*, *Nvvasa2* and *Nvnanos2* are expressed solely in the mesenterial tissues, excluded from the epithelia of the body wall and interestingly, are not detected within the physa of juveniles [86]. Those data combined with our current work further strengthen the idea that a specific stem cell population, able to respond to the amputation stress is located in the mesenteries of *Nematostella*.

However, it is important to note that the germ line of anthozoans is located in the mesenteries [86]. This of course raises the questions if i) the germ line cells correspond to the pool of pluripotent stem cells involved in the regeneration process, or if ii) the mesenteries contain in addition to the germ line, a pool of set aside stem cells that respond to the amputation stress by migrating towards the wound site to reform missing body parts. If the first idea would be true, then one would expect that a tradeoff between sexual reproduction and regeneration might exist, as regeneration would exhaust the germ-line pool for the tissue repair process. First experiments to analyze if sexual reproduction is impaired after bisection, revealed that the spawning capacity of the regenerated adult polyps remained unchanged (data not shown). While this suggests that the germ line is different from the stem cell population in the mesenteries, additional experiments are required to fully address this question.

Taken together, we provide a list of evidences that strongly support the idea that there are at least two populations of stem cells in *Nematostella* that are actively deployed and act in synergy during the regeneration process. However, further cellular (self-renewal, potency) as well as molecular characterization (e.g. stem cell/pluripotency markers) is needed in order to characterize in details their stem cell identity.

### The presence of mesenteries as specific feature of anthozoans may account for a spatial dissociation of the signal-emitting and signal-receiving compartments required for regeneration

Hydrozoans lack mesenteries and their pluripotent stem cells (i-cells) are located in the epithelia of the midgastric region and are able to give rise to the germ line as well as somatic cells [87–89]. The present study revealed that anthozoans possess stem cell populations that are located throughout the body within the epithelia but importantly also within the mesenteries. Because the oocytes (germ line) are located within the mesenteries, it is highly probable that, if any pluripotent stem cells (giving rise to germ and soma) exist in *Nematostella* they are located within the mesenteries. Taken into an evolutionary context one proposition would be that hydrozoans (*Hydra, Hydractinia*) cope the lack of mesenteries with the existence of pluripotent stem cells (at the origin of germ and somatic cells) as well as a regeneration-inducing signal localized both directly within the epithelia. Our present results from the anthozoan sea anemone *Nematostella vectensis* show that the regeneration signal emitting (mesenteries) and receiving (epithelia) tissues are spatially separated and that a synergic effect of the stem cell populations is required to complete the regeneration process. We thus propose that the emergence/loss of structure complexity/compartmentalization can influence the proprieties of tissue plasticity, changes the competence of tissues to reprogram and, in the context of regeneration, the capacity to emit or respond to a regeneration-inducing signal.

## Conclusions

Although cnidarian regeneration has intrigued scientists for over 300 years, little was known about the origin of the regeneration inductive signal, the interplay between tissues and its role on coordinating a particular cellular dynamics required for a successful regeneration process. For the present study, we used *Nematostella vectensis,* an anthozoan cnidarian that possesses promising features to become a potent and complementary novel whole body regeneration model (reviewed in [44, 90]). By combining dissection and grafting experiments with functional and modern cellular approaches we have identified i) the body structure that emits a regeneration inductive signal, ii) the crucial importance of a tissue crosstalk to induce a cellular response and iii) that two populations of stem cells, activated by the amputation stress and tissue crosstalk, are synergically required to successfully complete the reformation of lost body parts. In addition to the identification of the previously unknown anthozoan stem cell populations, the tissue crosstalk we describe here has never been reported from other cnidarians. This is mainly due to differences observed in cnidarian anatomies and opens new opportunities to understand how the emergence/loss of structure complexity/compartmentalization can influence the proprieties of the tissue plasticity that compose the body of an animal. This in turn may have important implications on the competence of a tissue to reprogram and, in the context of regeneration, the capacity to emit or respond to a regeneration-inducing signal.

## Acknowledgements

The authors thank the current team members as well as the Saccani, Liti, Shkreli and Féral labs for stimulating discussions on the project. We are grateful to Marina Shkreli for careful reading and comments on the manuscript. We also thank Isabelle Bourget for running the irradiation device and the Pasteur-IRCAN Molecular and Cellular Imaging Core Facility (PICMI) for providing access to the Zeiss LSM Exciter confocal microscope. PICMI was supported financially by: le Cancéropole PACA, la Région Provence Alpes-Côte d’Azur, le Conseil Départemental 06 and l’INSERM.

## Availability of data and materials

All biological material presented in this study is available upon request.

## Authors’ contributions

ARA and ER conceived and designed experiments. ARA, KF and SF performed experiments, generated, collected and analyzed data. ER contributed reagents, materials, and analysis tools. ARA and ER drafted the manuscript. All authors read and approved the final manuscript.

## Additional Files: Figure legends

**Figure S1.**
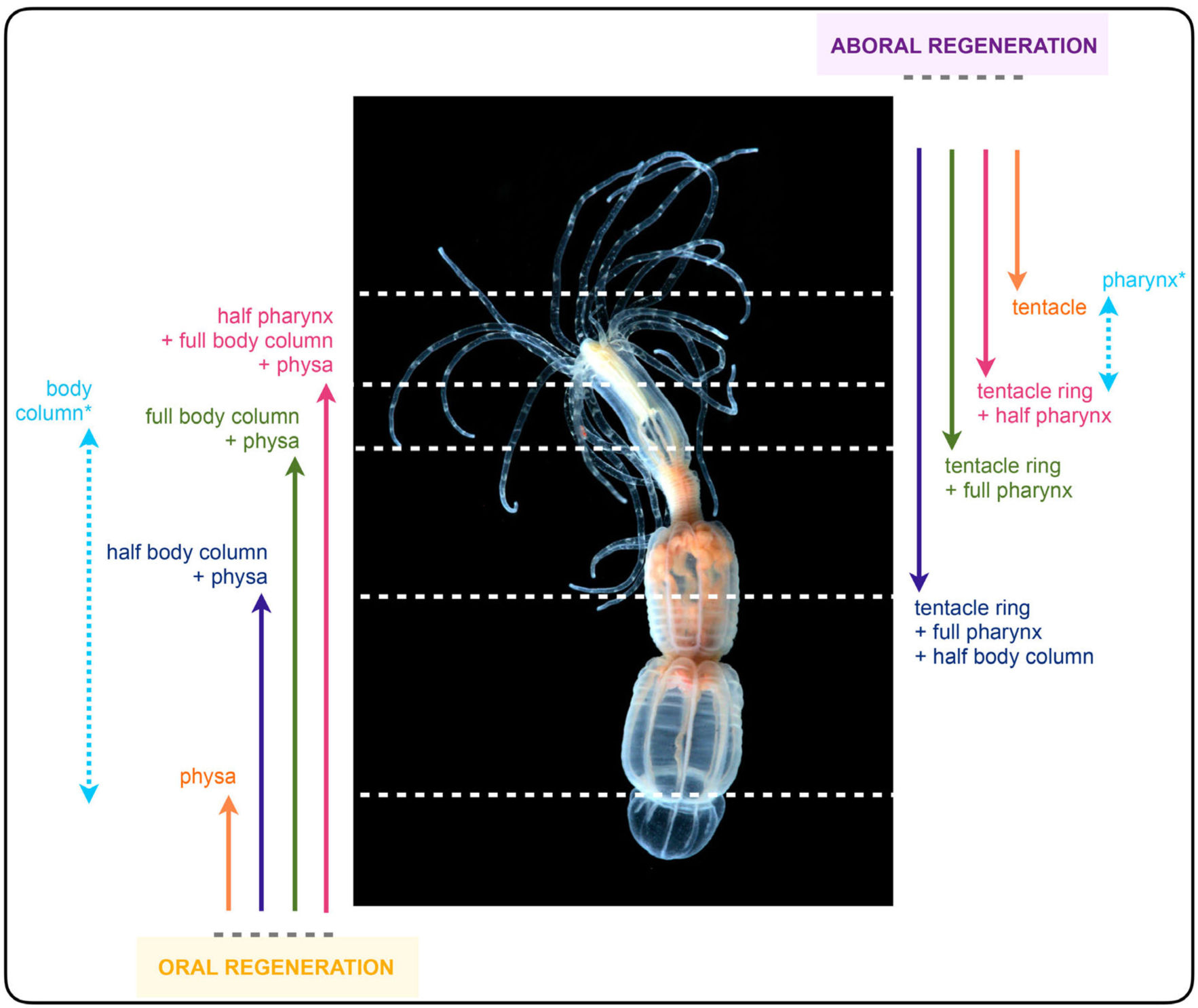
Diagram summarizing the various amputation sites along the body column of *Nematostella* used in this study. The asterisks for [body column] and [pharynx] indicate that in those isolated parts of the animal, oral and aboral regeneration were scored.

**Figure S2.**
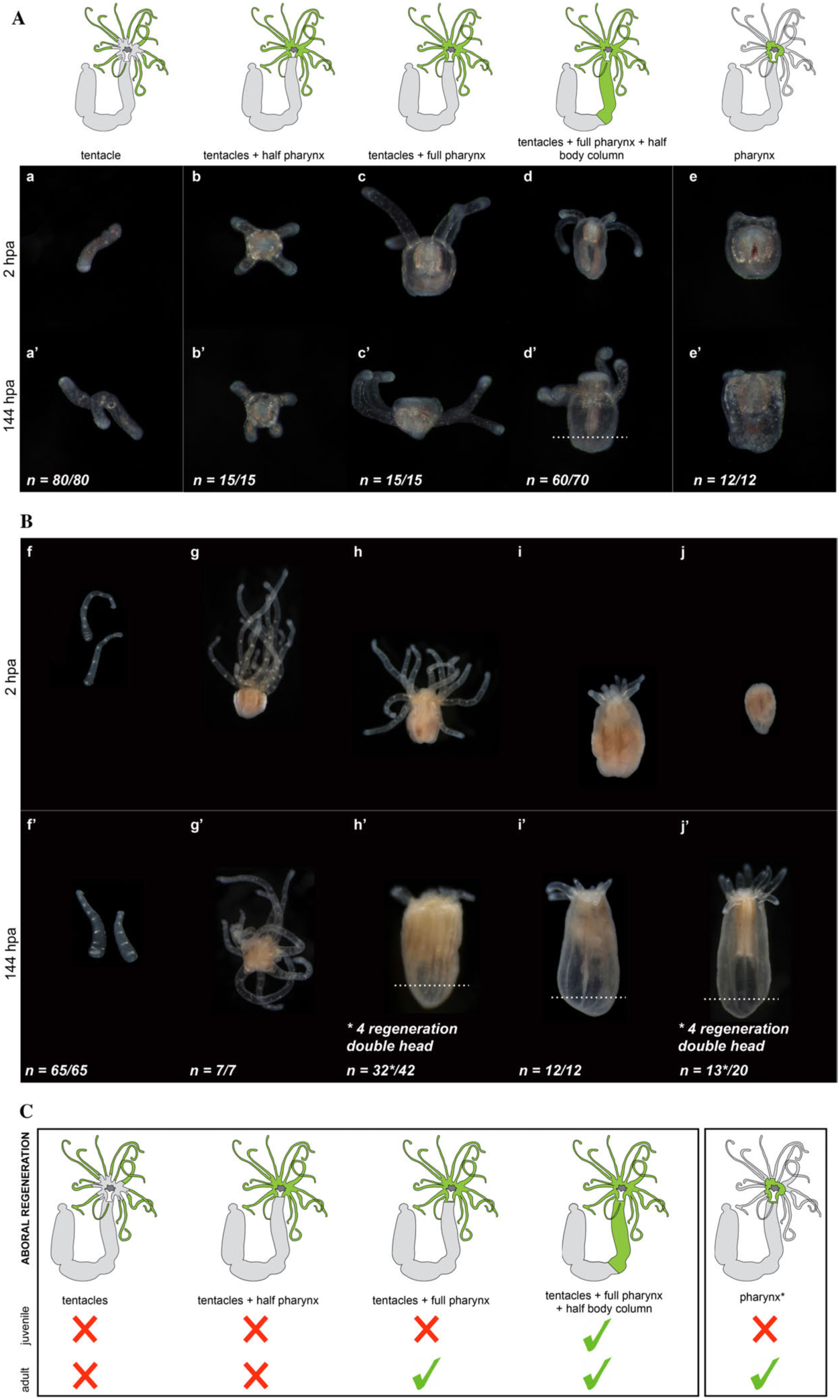
Aboral regenerative capacity analyzed in juveniles (Aa-e, Aa’-e’) and in adults (Bf-j, Bf’-j’) from isolated [tentacles] (Aa, Aa’, Bf, Bf’); [tentacles + half pharynx] (Ab, Ab’, Bg, Bg’); [tentacles + full pharynx] (Ac, Ac’, Bh, Bh’); [tentacles + half pharynx + half body column] (Ad, Ad’, Bi, Bi’); [pharynx] (Ae, Ae’, Bj, Bj’). The success of aboral regeneration is indicated by the development of a new physa delimitated by the white dashed line. Photographs of the isolated body parts at 2hpa (Aa-e, Bf-j). Phenotypes observed after 144hpa (6 days post amputation) (Aa’-e’, Bf’-j’). n=[number of specimen with represented phenotype]/[total number of analyzed specimen]. (*) indicates that from the pool of isolated body parts, four animals regenerated with two oral but no aboral regions. **C**. Diagram summarizing the aboral regeneration experiments carried out in juveniles and adults. Green parts in the schematic *Nematostella* indicate the isolated body part, green checkmarks the regenerative success and red crosses the absence of regeneration after body part isolation.

**Figure S3.**
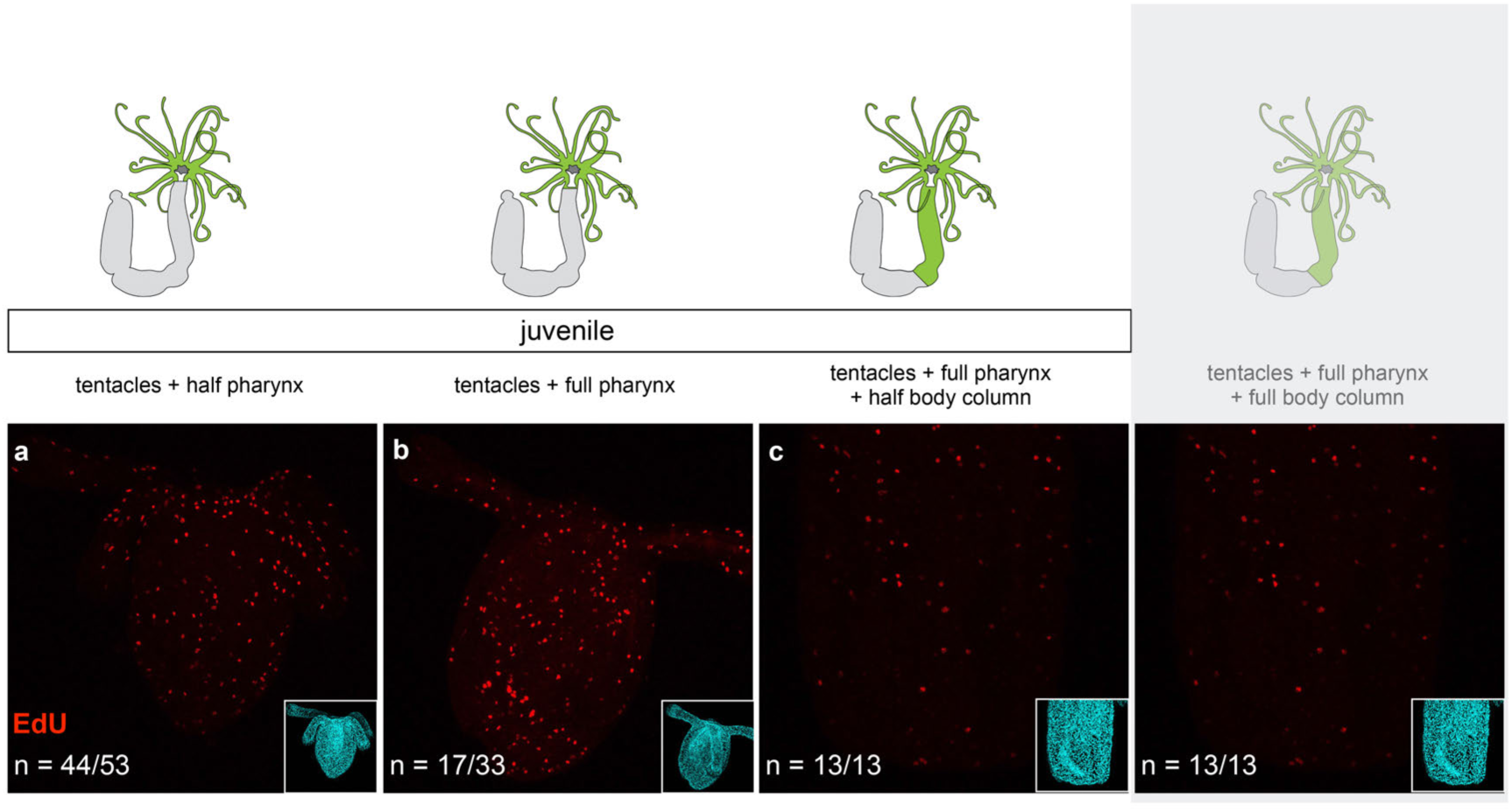
Cell proliferation at 48hpa in [tentacles + half pharynx] (a); [tentacles + full pharynx] (b); [tentacles + full pharynx + half body column] (c). Confocal microscopy images showing cellular proliferation (EdU labeling, a-c; red) and nuclei using DAPI (cyan, right bottom square) for corresponding images. n=[number of specimen with represented phenotype]/[total number of analyzed specimen].

**Figure S4.**
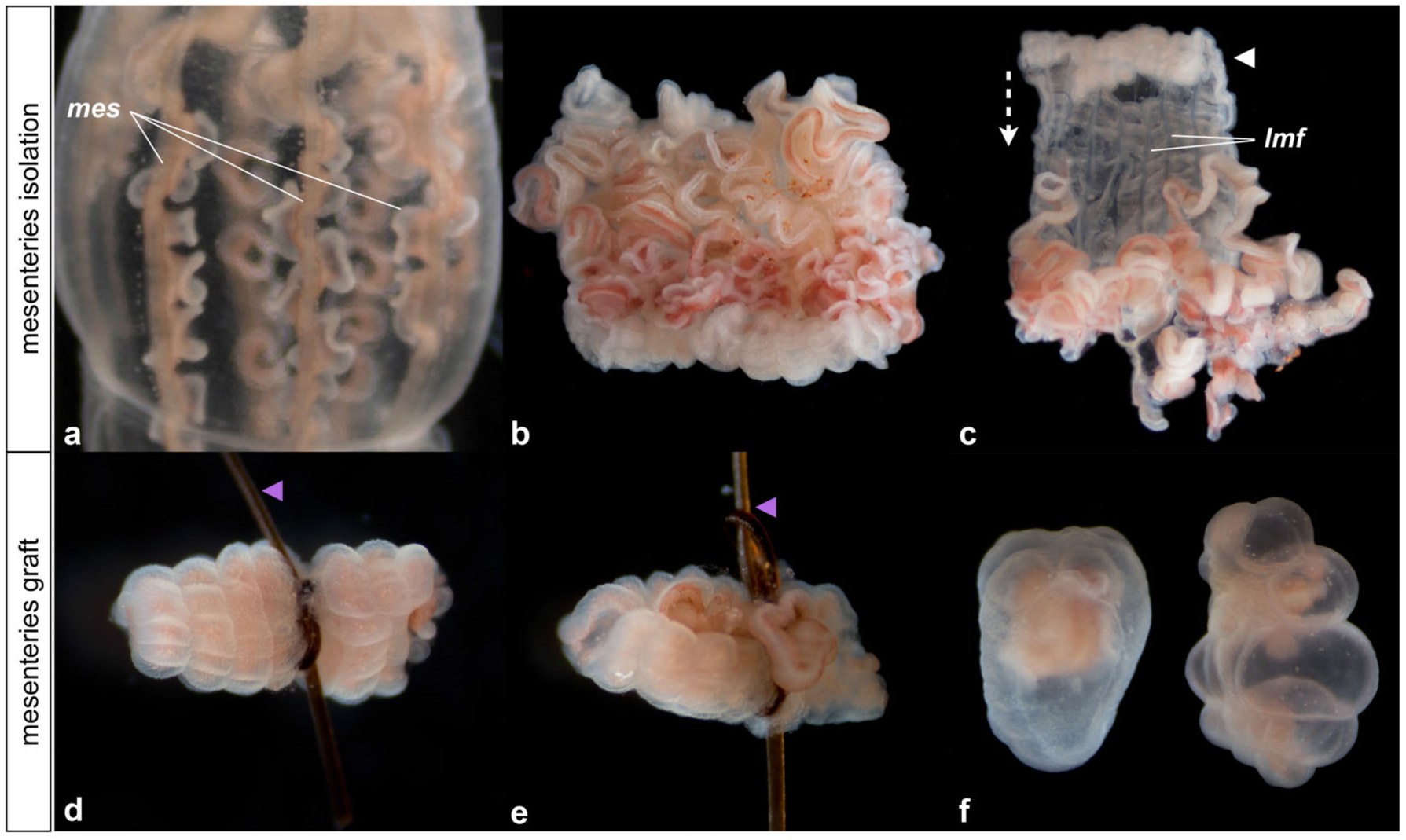
Macrophotographs illustrating the tissue isolation and grafting experiments. (a-c) are images of the experimental protocol for the isolation of the mesenteries and the body wall epithelia. (d, e) are images of the [mes] + [ecto + gastro] grafts immediately after the combination. Purple arrowheads indicate the organic string required to maintain the tissues together. (f) is an image of two samples three days after grafting the tissues together and in which the graft between the mesenteries and the body wall epithelia was successful (and later regenerated). Note that the integrity of the mesenterial tissue (pink) is maintained.

**Figure S5.**
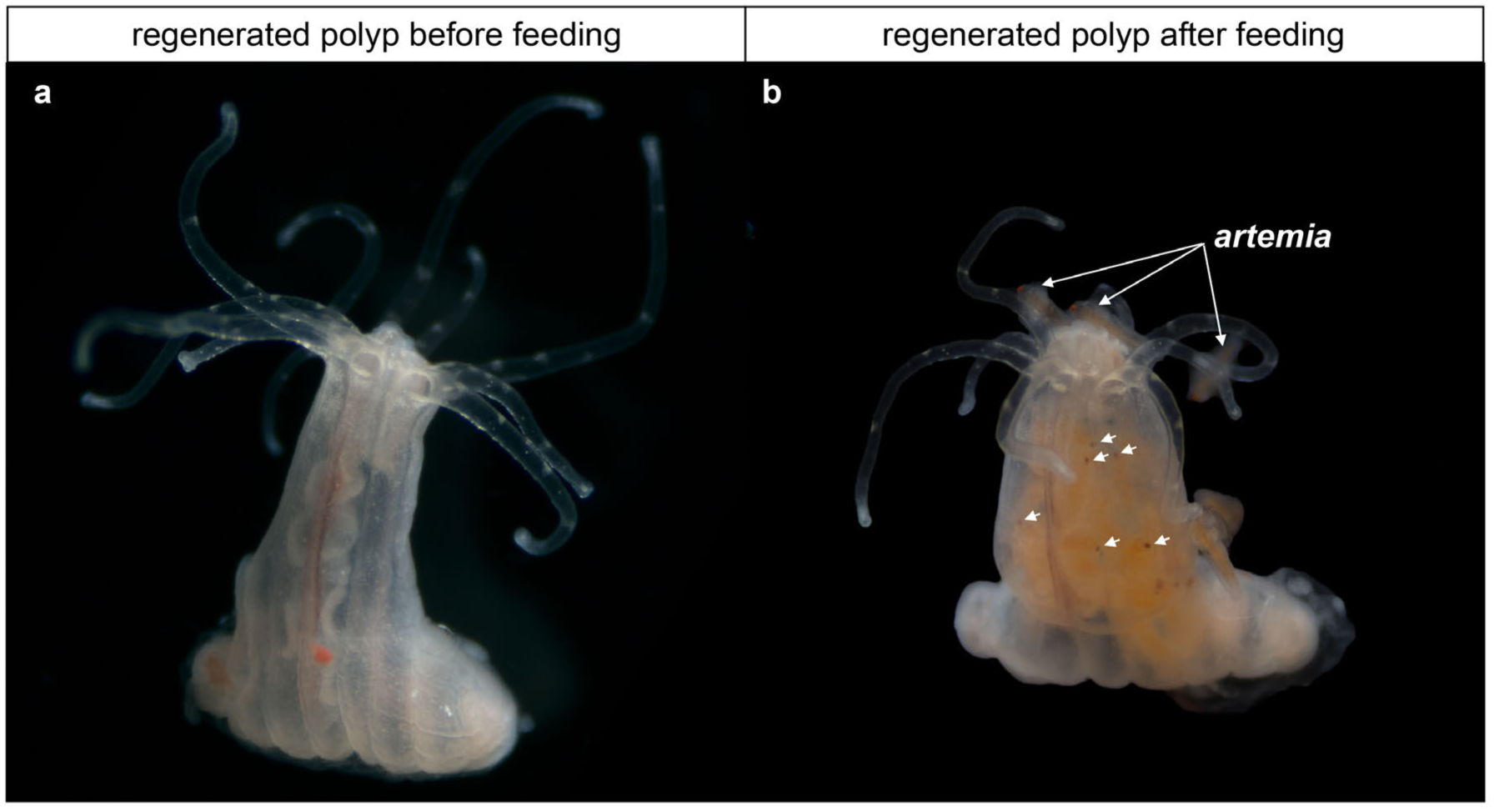
Macrophotographs illustrating regenerated polyps before feeding (a) and after feeding (b). The white arrows in (b) show the eaten artemia (brine shrimp).

**Figure S6.**
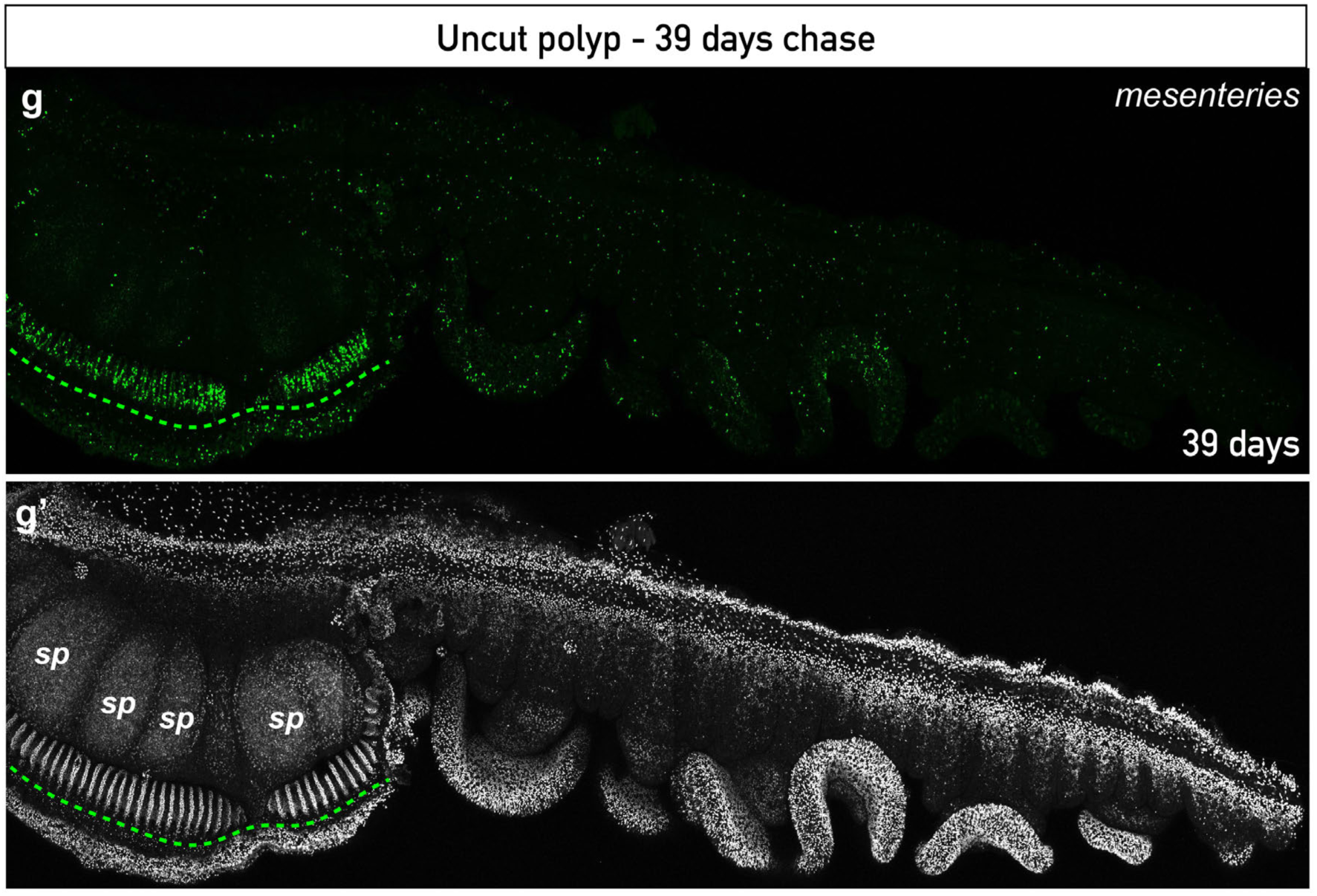
Presence of Label (EdU) Retaining cells (LRCs) in uncut adult gonads. The discontinued line (g,g’) show the LRCs and the zone of LRCs accumulation in the gonad, respectively. LRC EdU staining in green (g) and DAPI staining that is labeling the nuclei in white (g’). *sp*, sperm mass.

**Figure S7.**
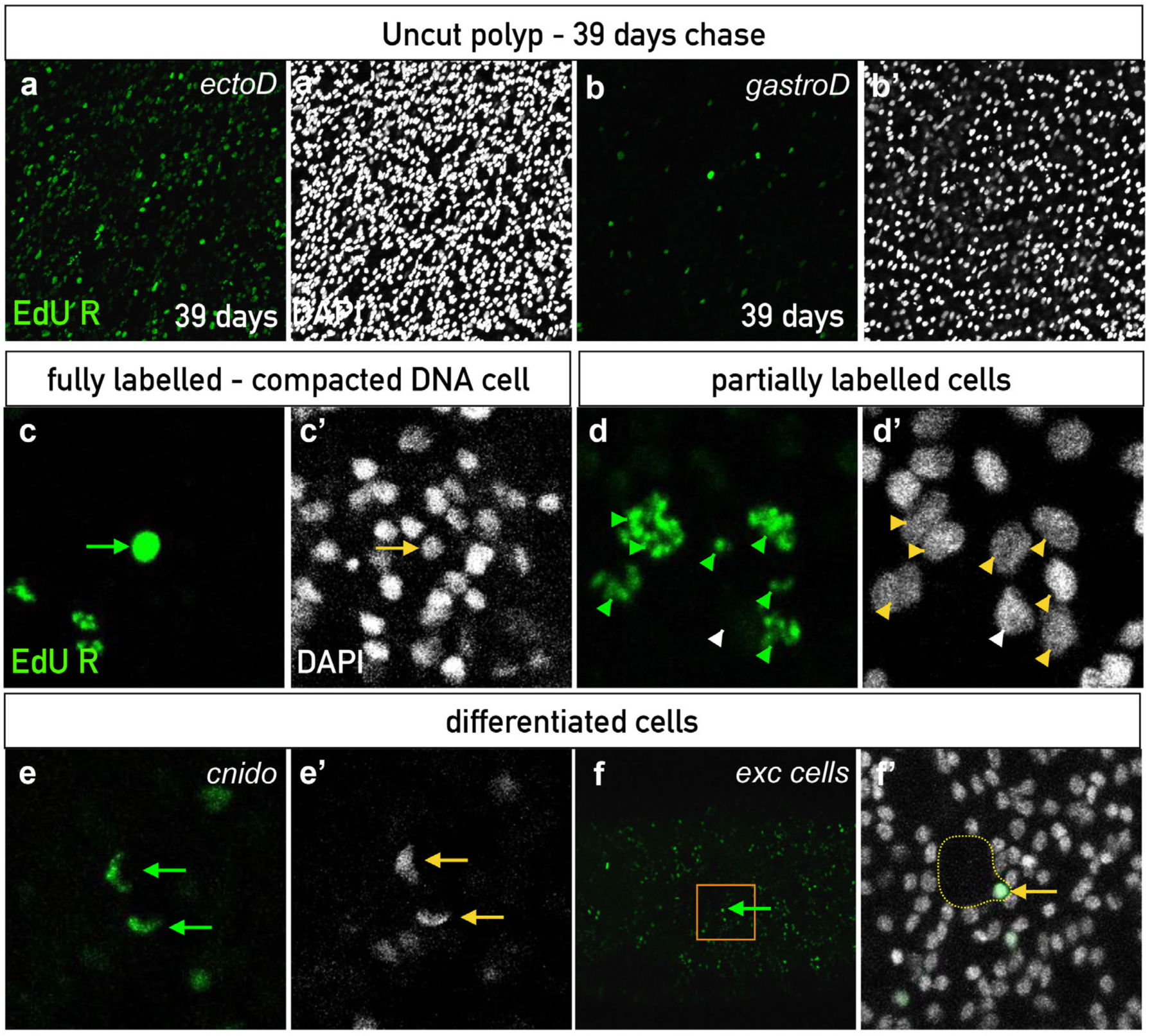
Presence of Label (EdU) Retaining cells (LRCs) in uncut adults. Confocal images of the LRCs in adult epithelia chased at 39 days (EdU R in green, a-e) after a one-week pulse. Nuclei are in white (DAPI) (a’-e’). Overlay of EdU and DAPI staining in d’. The renewal of the gastrodermal cells seems faster than the ectodermal cells as LRCs are more isolated and sparse in the gastrodermis than in the ectodermis in adults (a,b). Within the LRCs, one can distinguish differentiated cells such as cnidocytes (e,e’), excretory cells (f,f’; discontinued line in f’) and batteries of nematocysts. The latter have partially labeled nuclei potentially because of their several rounds of cell division (d,d’). Cells with highly condensed DNA are also found (c,c’). Green arrows (c,e-f) and arrow heads (d) indicate the LRCs EdU staining. Yellow arrows (c’,e’-f’) and arrow heads (d’) indcate the LRCs nuclei. Note that one cell of the batteries of the nematocysts is poorly or not labeled (white arrow head in d’) probably due to the dilution of the signal through cell divisions. f’ is a magnification of the square represented in f. *ectoD,* ectodermal view; *exc cells*, excretory cells; *gastroD*, gastrodermal view.

**Figure S8.**
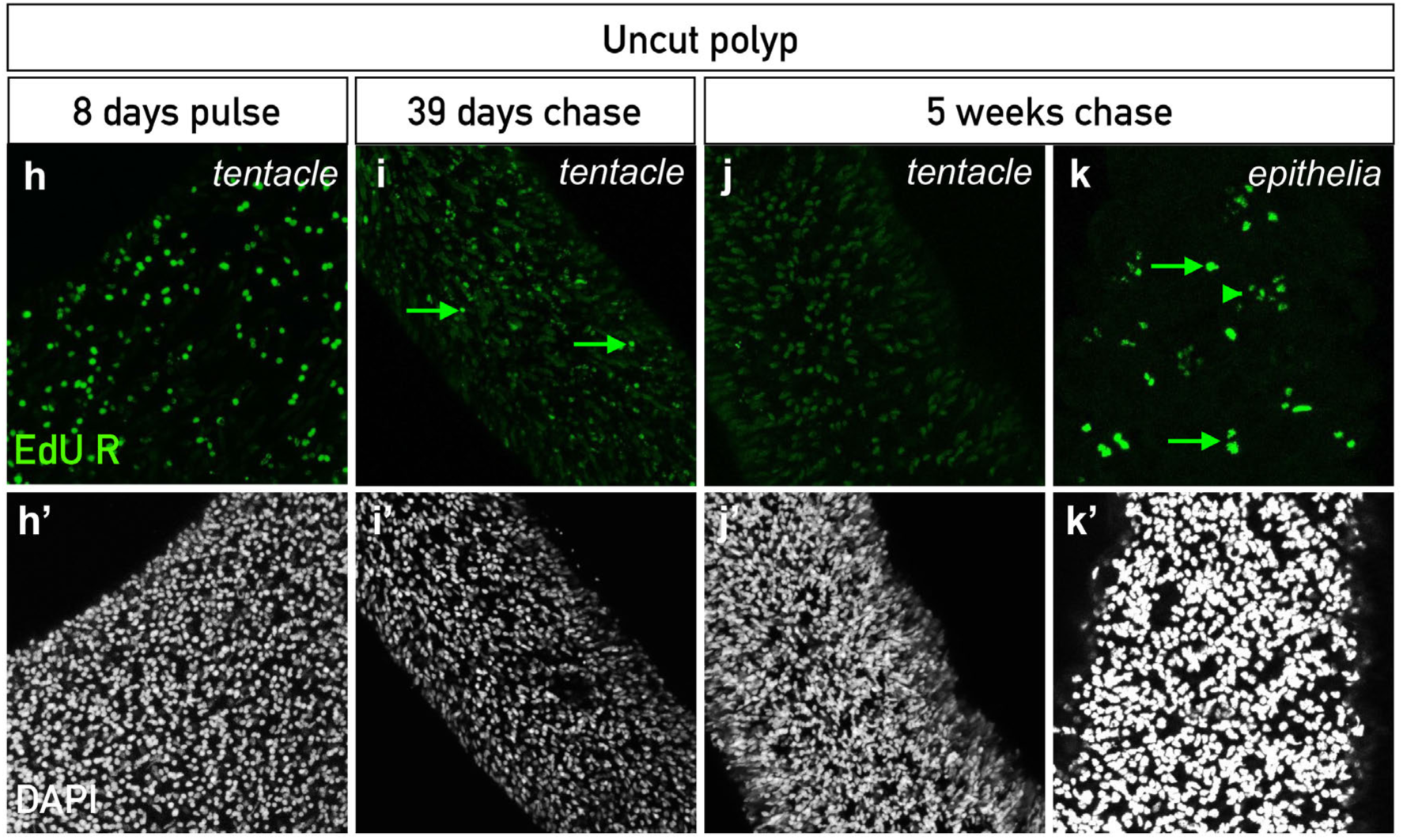
LRCs at 8 days after pulse (h), 39 days chase (i) and 5 months chase (j) in the tentacles of the uncut polyp. LRCs are present in the tentacles up to 39 days (i) but not at 5 months chase (i). However, they are still present in the body wall epithelia of the 5 months chase (k). LRC EdU staining in green (h-k) and DAPI staining labeling the nuclei in white (h’-k’). Green arrows (h,i,k) show the LRCs. Green arrow-head in k shows potential batteries of nematocysts.

**Figure S9.**
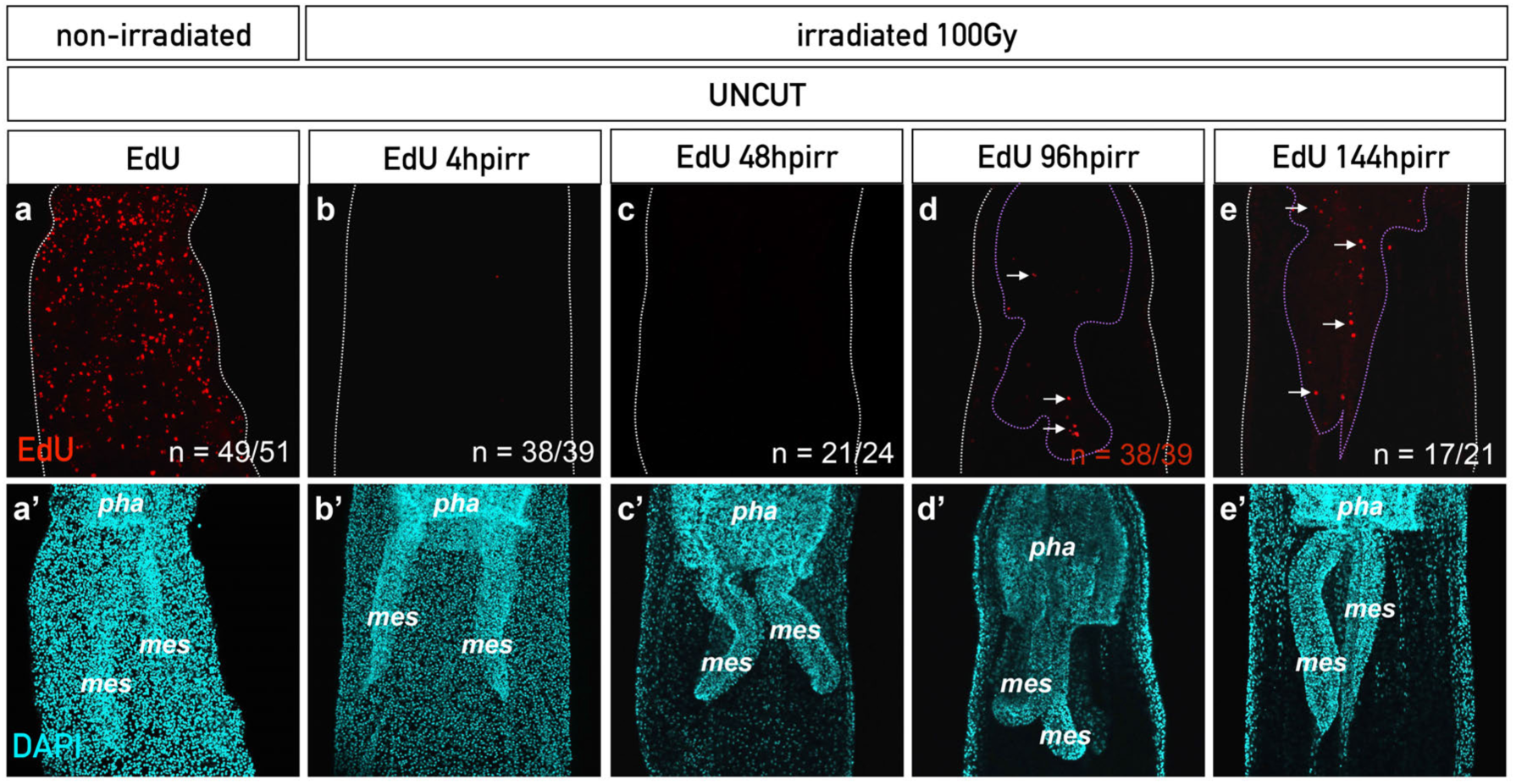
Irradiation blocks cell proliferation (a,a’,b,b’) and a population of cells is able to re-inter the mitotic cycle between 48 and 96 hours post irradiation (hpirr) in the uncut animal. Confocal images of control non irradiated (a,a’) and irradiated (b-e,b’-e’) uncut polyps. While cell proliferation is blocked after irradiation (a,a’,b,b’), a few cells in the mesenteries, were able to re-enter the cell cycle between 48 and 96hpirr. They were also detectable at 144hpirr within the mesenteries and in the pharyngeal region (e,e’). *mes*, mesenteries; *pha*, pharynx. n=[number of specimen with represented phenotype]/[total number of analyzed specimen].

**Figure S10.**
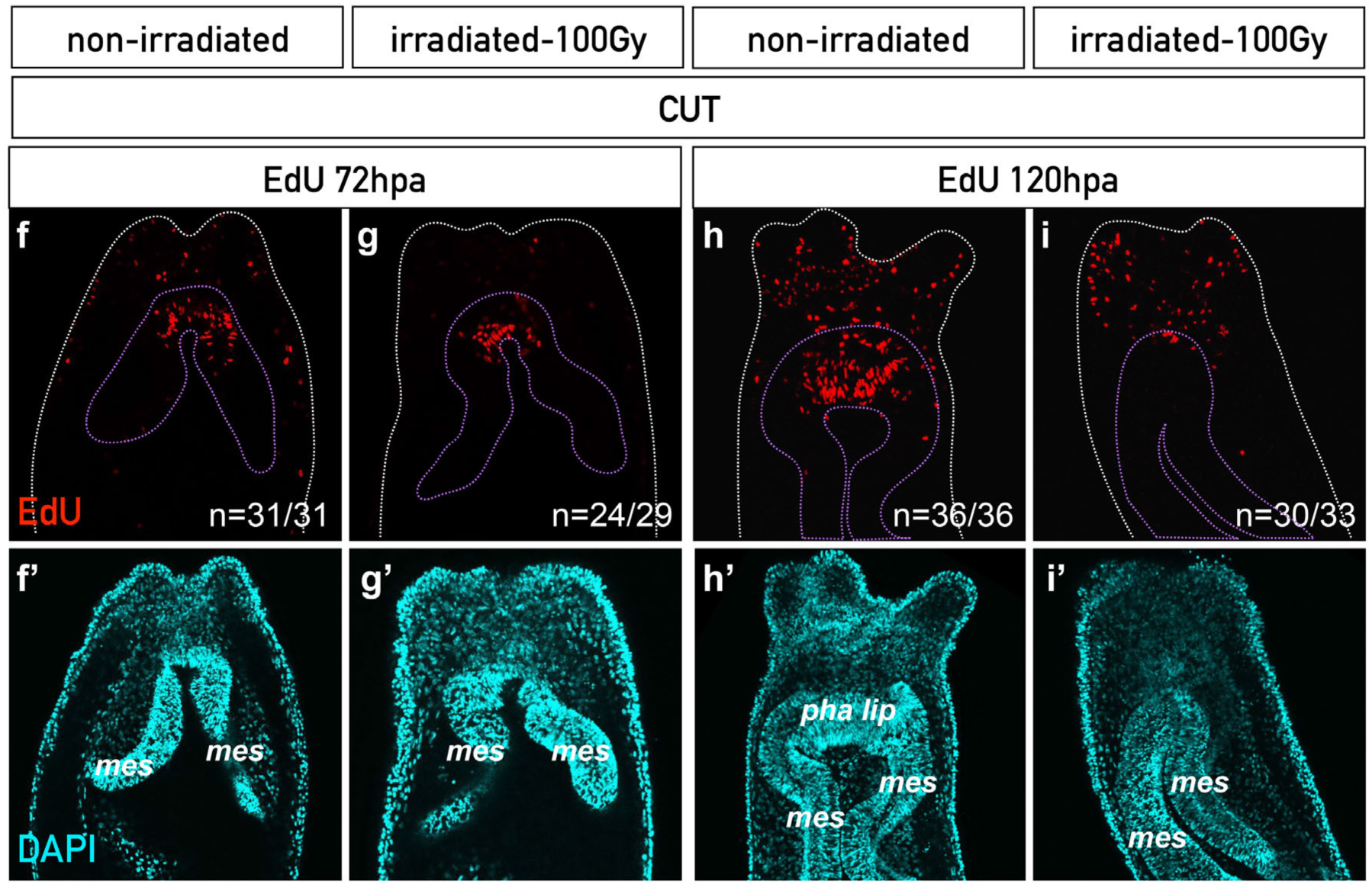
A population of cells is able to re-enter the mitotic cycle between 24 and 48 hours post amputation (hpa) (corresponding to 28 and 52hpirr) in the 100 Grays irradiated polyps. Confocal images of control non irradiated (f,f’: 72hpa; h,h’: 120hpa) and irradiated (g,g’: 72hpa; i,i’: 120hpa) polyps. The cells in the mesenteries, localized at the amputation site, that re-entered the cell cycle between 24 and 48hpa are also detectable at 72 and 120hpa (g,i). EdU staining in red shows cell proliferation (f,i). Nuclei are in cyan (DAPI) (f’,i’). *mes*, mesenteries; *pha lip*, pharyngeal lip; *ten*, tentacles. n=[number of specimen with represented phenotype]/[total number of analyzed specimen].

**Figure S11.**
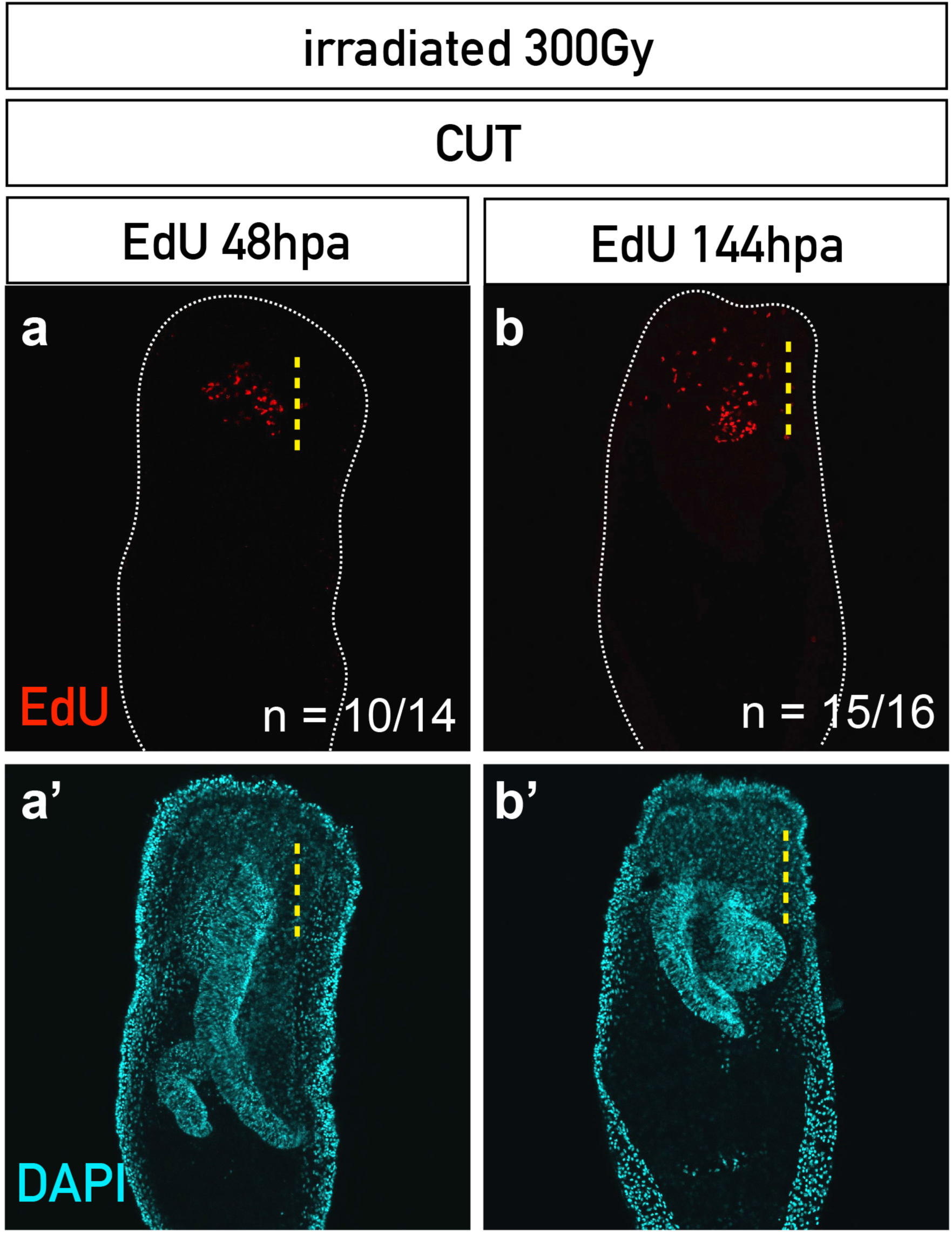
A population of cells is able to re-enter the mitotic cycle between 24 and 48 hours post amputation (hpa) (corresponding to 28 and 52hpirr) in 300 Grays irradiated polyps. Confocal images of irradiated and amputated polyp (j,j’: 48hpa; k,k’: 144hpa). EdU staining in red shows cell proliferation (j,k). Nuclei are in cyan (DAPI) (j’,k’). *mes*, mesenteries. n=[number of specimen with represented phenotype]/[total number of analyzed specimen].

**Figure S12.**
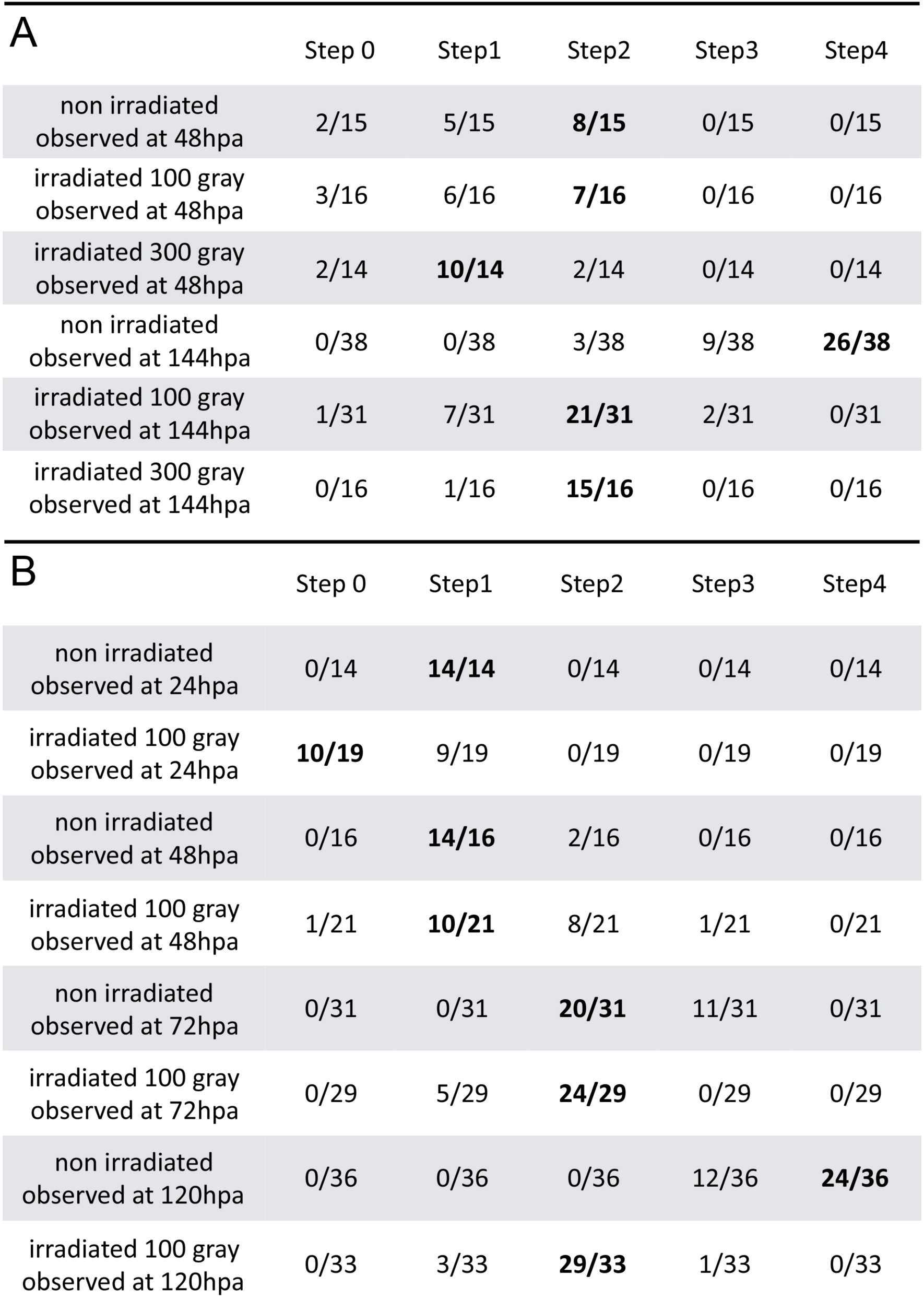
The resulting phenotype from non-irradiated *vs* irradiated uncut animals that were bisected and then scored for regeneration progression at various time points after amputation in two independent experiments (**A**: experiment batch 1 and **B**: experiment batch 2).

**Figure S13.**
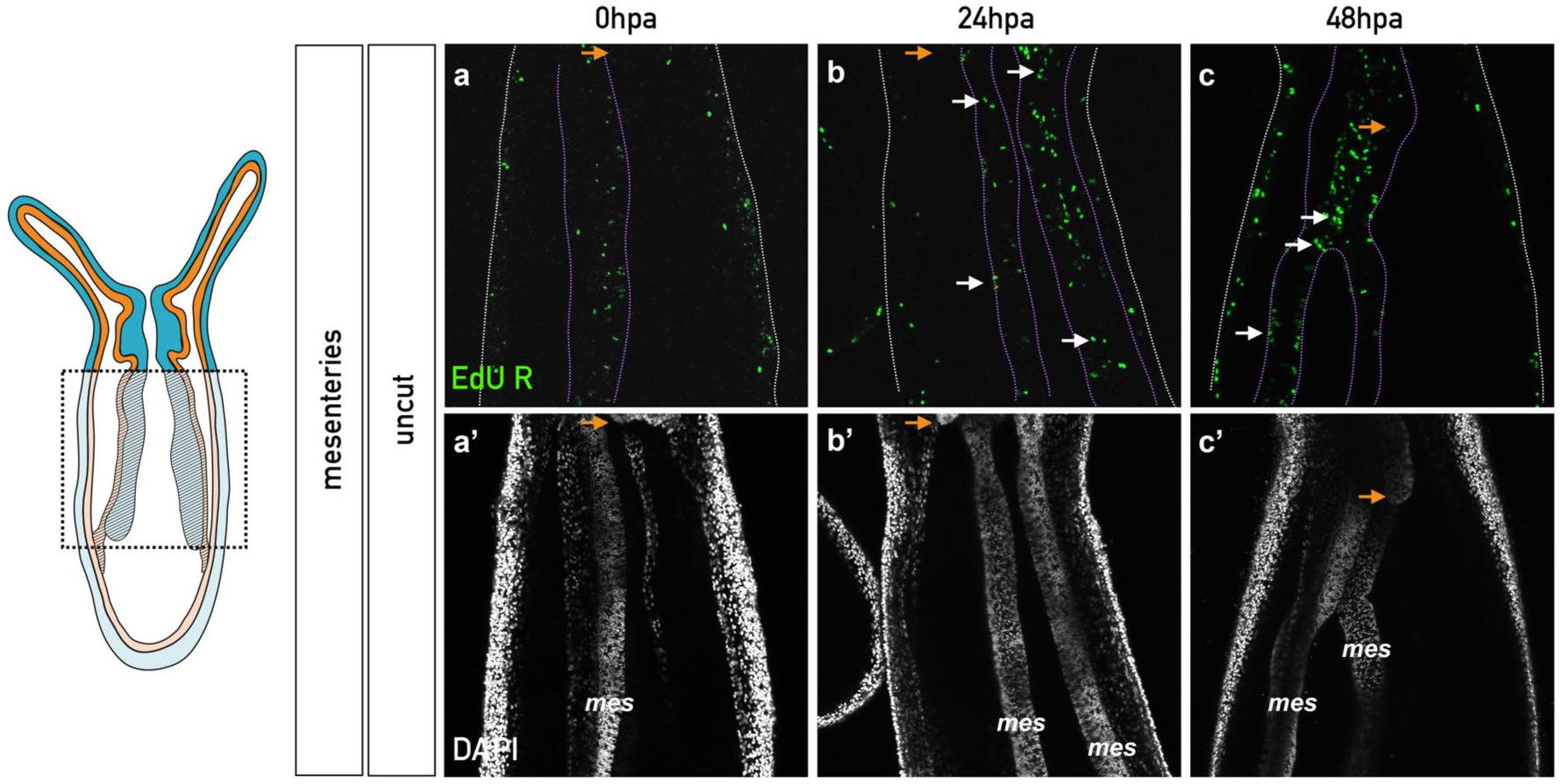
Chase of the EdU+ cells at 0, 24 and 48 hours post-1hour-pulse (hpp) in control uncut animals. Confocal images of EdU+ cells (EdU R) that are stained in green (a-c) and the nucleus in white (a’-c’). Orange arrows indicate the position of the pharynx. White arrows indicate EdU R cell pairs (b,c). No accumulation of EdU+ retaining cells is detected in uncut animals (Aa-d). The discontinued white and purple lines indicate the contours of the epithelia and mesenteries, respectively. *mes*, mesenteries.

**Figure S14.**
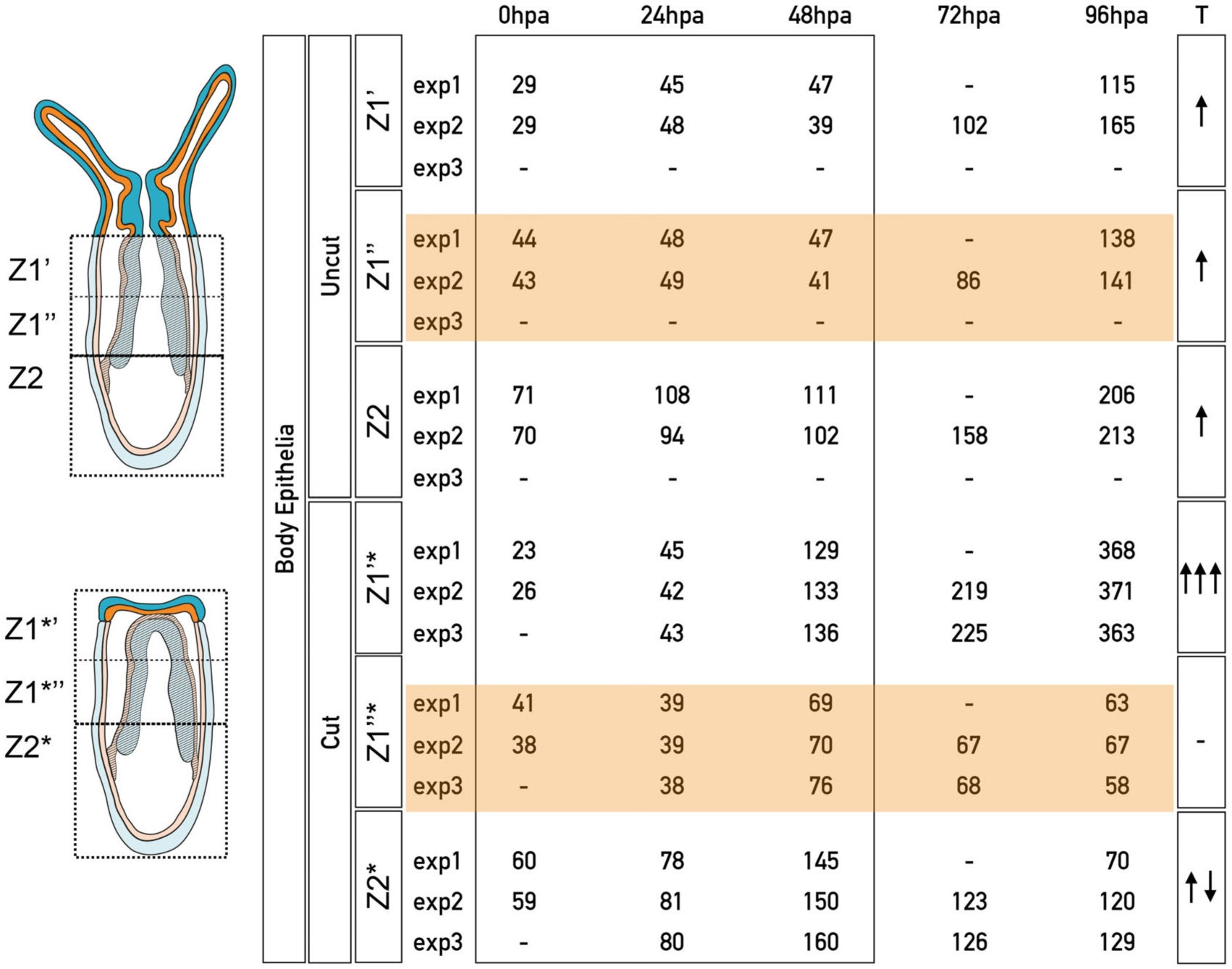
Average number of EdU R cells during the time course of regeneration in three different zones located along the oral-aboral axis of the body epithelia for uncut and cut (*) polyps. These zones are named Z1’, Z1’’ and Z2. All the described zones are restricted to the epithelia; mesenteries are excluded from the counting. The zones are defined as followed: Z1 - oral half of the polyp; Note that in the uncut animal, the region above the sub-pharyngeal line (the head) was not taken into account to define the zone; Z1’ and Z1’’ are the oral and aboral halves of Z1, respectively. Z2 - aboral half of the polyp. (*) is associated to the three zones defined in regenerating polyps (Z1’*, Z1’’* and Z2*). Counting was performed in three independent experiments (*exp1, exp2, exp3*). Arrows on the right side indicate general variations of the average number of cells for the different zones in the uncut *vs* cut animals.

## References

1. King RS, Newmark PA. Beyond the cell: The cell biology of regeneration. The Journal of Cell Biology. 2012;196:553–62.

2. Trembley A. Mémoires pour servir à l’histoire d‘un genre de polypes d’eau douce, à bras en forme de cornes. Verbeek JH, editor. Leiden; 1744;:1–404.

3. Réaumur RA. Animaux coupés et partagés en plusieurs parties, et qui se reproduisent tout entiers dans chacune. (null), editor. Memoires de lAcademie Royale des Sciences de Paris. 1741;:33–5.

4. Sánchez Alvarado A, Tsonis PA. Bridging the regeneration gap: genetic insights from diverse animal models. Nat Rev Genet. 2006;7:873–84.

5. Galliot B, Schmid V. Cnidarians as a model system for understanding evolution and regeneration. Int J Dev Biol. 2002;46:39–48.

6. Holstein TW, Hobmayer E, Technau U. Cnidarians: an evolutionarily conserved model system for regeneration? Dev Dyn. 2003;226:257–67.

7. Bely AE, Nyberg KG. Evolution of animal regeneration: re-emergence of a field. Trends Ecol Evol (Amst). 2010;25:161–70.

8. Pallas PS. **Fasciola Punctata.** Spicilegia zoologica - Quibus novae imprimis et obscurae animalium species iconibus, descriptionibus atque comentariis illustrantur. 1774;:22–3.

9. Morgan TH. Experimental studies of the regeneration of Planaria maculata. Archiv für Entwickelungsmechanik der Organismen. Springer-Verlag; 1898;7:364–97.

10. Reddien PW, Alvarado AS. Fundamentals of planarian regeneration. Annu Rev Cell Dev Biol. 2004;20:725–57.

11. Wenemoser D, Reddien PW. Planarian regeneration involves distinct stem cell responses to wounds and tissue absence. Dev Biol. 2010;344:979–91.

12. David CN, Plotnick I. Distribution of interstitial stem cells in Hydra. Dev Biol. 1980;76:175–84.

13. Bode HR. Head regeneration inHydra. Dev Dyn. 2003;226:225–36.

14. Hazen AP. Regeneration in Hydractinia and Podocoryne. American Naturalist. 1902.

15. Schmid V, Schmid B, Schneider B, Stidwill R, Baker G. Factors effecting manubrium-regeneration in hydromedusae (Coelenterata). Wilhelm Roux’ Archiv. Springer-Verlag; 1976;179:41–56.

16. Duffy DJ, Plickert G, Kuenzel T, Tilmann W, Frank U. Wnt signaling promotes oral but suppresses aboral structures in Hydractinia metamorphosis and regeneration. Development (Cambridge, England). 2010;137:3057–66.

17. Frank U, Leitz T, Müller WA. The hydroid Hydractinia: a versatile, informative cnidarian representative. Bioessays. John Wiley & Sons, Inc; 2001;23:963–71.

18. Marcum BA, Campbell RD. Development of Hydra lacking nerve and interstitial cells. J Cell Sci. 1978;29:17–33.

19. Galliot B. Regeneration in Hydra. eLS. Chichester, UK: John Wiley & Sons, Ltd; 2013.

20. Chera S, Ghila L, Dobretz K, Wenger Y, Bauer C, Buzgariu W, et al. Apoptotic Cells Provide an Unexpected Source of Wnt3 Signaling to Drive Hydra Head Regeneration. Dev Cell. Elsevier Ltd; 2009;17:279–89.

21. Gahan JM, Bradshaw B, Flici H, Frank U. The interstitial stem cells in Hydractinia and their role in regeneration. Curr Opin Genet Dev. 2016;40:65–73.

22. Bradshaw B, Thompson K, Frank U. Distinct mechanisms underlie oral versus aboral regeneration in the cnidarian Hydractinia echinata. eLife. 2015;4.

23. Müller WA, Teo R, Frank U. Totipotent migratory stem cells in a hydroid. Dev Biol. 2004;275:215–24.

24. Govindasamy N, Murthy S, Ghanekar Y. Slow-cycling stem cells in hydra contribute to head regeneration. Biology Open. 2014;3:1236–44.

25. Bridge D, Cunningham CW, Schierwater B, DeSalle R, Buss LW. Class-level relationships in the phylum Cnidaria: evidence from mitochondrial genome structure. Proceedings of the National Academy of Sciences. National Academy of Sciences; 1992;89:8750–3.

26. Kim J, Kim W, Cunningham CW. A new perspective on lower metazoan relationships from 18S rDNA sequences. Mol Biol Evol. 1999.

27. Medina M, Collins AG, Silberman JD, Sogin ML. Evaluating hypotheses of basal animal phylogeny using complete sequences of large and small subunit rRNA. Proceedings of the National Academy of Sciences. National Acad Sciences; 2001;98:9707–12.

28. Collins AG, Schuchert P, Marques AC, Jankowski T, Medina M, Schierwater B. Medusozoan phylogeny and character evolution clarified by new large and small subunit rDNA data and an assessment of the utility of phylogenetic mixture models. Syst. Biol. Oxford University Press; 2006;55:97–115.

29. Dunn CW, Hejnol A, Matus DQ, Pang K, Browne WE, Smith SA, et al. Broad phylogenomic sampling improves resolution of the animal tree of life. Nature. Nature Publishing Group; 2008;452:745–9.

30. Philippe H, Derelle R, Lopez P, Pick K, Borchiellini C, Boury-Esnault N, et al. Phylogenomics revives traditional views on deep animal relationships. Curr Biol. 2009;19:706–12.

31. Erwin DH. Wonderful Ediacarans, wonderful cnidarians? Evol Dev. Blackwell Publishing Inc; 2008;10:263–4.

32. Gold DA, Jacobs DK. Stem cell dynamics in Cnidaria: are there unifying principles? Dev Genes Evol. 2012;223:53–66.

33. Zapata F, Goetz FE, Smith SA, Howison M, Siebert S, Church SH, et al. Phylogenomic Analyses Support Traditional Relationships within Cnidaria. Steele RE, editor. PLoS ONE. Public Library of Science; 2015;10:e0139068.

34. Daly M, Brugler MR, Cartwright P, Collins AG. The phylum Cnidaria: a review of phylogenetic patterns and diversity 300 years after Linnaeus. 2007.

35. Zhang ZQ. Animal biodiversity: An introduction to higher-level classification and taxonomic richness. Zootaxa. 2011.

36. Fautin D, Mariscal R. Microscopic anatomy of invertebrates: Anthozoa. Harrison FW, Ruppert EE, editors. 1991;:1–93.

37. Won J, Rho B, Song J. A phylogenetic study of the Anthozoa (phylum Cnidaria) based on morphological and molecular characters. Coral reefs. Springer-Verlag; 2001;20:39–50.

38. Babonis LS, Martindale MQ, Ryan JF. Do novel genes drive morphological novelty? An investigation of the nematosomes in the sea anemone Nematostella vectensis. BMC Evol Biol. BMC Evolutionary Biology; 2016;16:1–22.

39. Hand C, Uhlinger KR. The culture, sexual and asexual reproduction, and growth of the sea anemone Nematostella vectensis. Biol Bull. MBL; 1992;182:169–76.

40. Fritzenwanker JH, Technau U. Induction of gametogenesis in the basal cnidarian Nematostella vectensis (Anthozoa). Dev Genes Evol. 2002;212:99–103.

41. Stefanik DJ, Friedman LE, Finnerty JR. Collecting, rearing, spawning and inducing regeneration of the starlet sea anemone, Nematostella vectensis. Nat Protoc. 2013;8:916– 23.

42. Putnam NH, Srivastava M, Hellsten U, Dirks B, Chapman J, Salamov A, et al. Sea anemone genome reveals ancestral eumetazoan gene repertoire and genomic organization. Science. 2007;317:86–94.

43. Layden MJ, Röttinger E, Wolenski FS, Gilmore TD, Martindale MQ. Microinjection of mRNA or morpholinos for reverse genetic analysis in the starlet sea anemone, Nematostella vectensis. Nat Protoc. 2013;8:924–34.

44. Layden MJ, Rentzsch F, Röttinger E. The rise of the starlet sea anemone Nematostella vectensis as a model system to investigate development and regeneration. WIREs Dev Biol. John Wiley & Sons, Inc; 2016.

45. Ikmi A, McKinney SA, Delventhal KM, Gibson MC. TALEN and CRISPR/Cas9-mediated genome editing in the early-branching metazoan Nematostella vectensis. Nature Communications. 2014;5:5486.

46. Kusserow A, Pang K, Sturm C, Lentfer J, Schmidt HA, Technau U, et al. Unexpected complexity of the Wnt gene family in a sea anemone. Nature. 2005;433:156–60.

47. Schwaiger M, Schonauer A, Rendeiro AF, Pribitzer C, Schauer A, Gilles AF, et al. Evolutionary conservation of the eumetazoan gene regulatory landscape. Genome Research. 2014.

48. Rentzsch F, Fritzenwanker JH, Scholz CB, Technau U. FGF signalling controls formation of the apical sensory organ in the cnidarian Nematostella vectensis. Development. 2008;135:1761–9.

49. Saina M, Technau U. Characterization of myostatin/gdf8/11 in the starlet sea anemone Nematostella vectensis. J Exp Zool. 2009.

50. Layden MJ, Boekhout M, Martindale MQ. Nematostella vectensis achaete-scute homolog NvashA regulates embryonic ectodermal neurogenesis and represents an ancient component of the metazoan neural specification pathway. Development (Cambridge, England). 2012;139:1013–22.

51. Röttinger E, Dahlin P, Martindale MQ. A Framework for the Establishment of a Cnidarian Gene Regulatory Network for “Endomesoderm” Specification: The Inputs of ß-Catenin/TCF Signaling. Mullins MC, editor. PLoS Genet. 2012;8:e1003164.

52. Leclère L, Rentzsch F. RGM Regulates BMP-Mediated Secondary Axis Formation in the Sea Anemone Nematostella vectensis. CellReports. 2014;9:1921–30.

53. Layden MJ, Johnston H, Amiel AR, Havrilak J, Steinworth B, Chock T, et al. MAPK signaling is necessary for neurogenesis in Nematostella vectensis. BMC Biol. 2016;14:61.

54. Genikhovich G, Fried P, Prünster MM, Schinko JB, Gilles AF, Fredman D, et al. Axis Patterning by BMPs: Cnidarian Network Reveals Evolutionary Constraints. CellReports. 2015;10:1646–54.

55. Wikramanayake AH, Hong M, Lee PN, Pang K, Byrum CA, Bince JM, et al. An ancient role for nuclear beta-catenin in the evolution of axial polarity and germ layer segregation. Nature. 2003;426:446–50.

56. Burton PM, Finnerty JR. Conserved and novel gene expression between regeneration and asexual fission in Nematostella vectensis. Dev Genes Evol. 2009;219:79–87.

57. Tucker RP, Shibata B, Blankenship TN. Ultrastructure of the mesoglea of the sea anemone Nematostella vectensis (Edwardsiidae). Invertebrate Biology. 2011;130:11–24.

58. Trevino M, Stefanik DJ, Rodriguez R, Harmon S, Burton PM. Induction of canonical Wnt signaling by alsterpaullone is sufficient for oral tissue fate during regeneration and embryogenesis in Nematostella vectensis. Dev Dyn. 2011;240:2673–9.

59. Passamaneck YJ, Martindale MQ. Cell proliferation is necessary for the regeneration of oral structures in the anthozoan cnidarian Nematostella vectensis. BMC Dev Biol. BMC Developmental Biology; 2012;12:1–1.

60. Bossert PE, Dunn MP, Thomsen GH. A staging system for the regeneration of a polyp from the aboral physa of the anthozoan cnidarian *Nematostella vectensis*. Dev Dyn. 2013;242:1320–31.

61. Schaffer AA, Bazarsky M, Levy K, Chalifa-Caspi V, Gat U. A transcriptional time-course analysis of oral vs. aboral whole-body regeneration in the Sea anemone Nematostella vectensis. BMC Genomics. 2016;17:718.

62. Amiel AR, Johnston HT, Nedoncelle K, Warner JF, Ferreira S, Röttinger E. Characterization of Morphological and Cellular Events Underlying Oral Regeneration in the Sea Anemone, Nematostella vectensis. Int J Mol Sci. 2015;16:28449–71.

63. Ormestad M, Amiel A, Röttinger E. Ex-situ Macro Photography of Marine Life. Imaging Marine Life. Weinheim, Germany: Wiley-VCH Verlag GmbH & Co. KGaA; 2013. pp. 210–33.

64. Cheung TH, Rando TA. Molecular regulation of stem cell quiescence. Nat Rev Mol Cell Biol. 2013;14:329–40.

65. Bickenbach JR. Identification and behavior of label-retaining cells in oral mucosa and skin. J. Dent. Res. 1981;60 Spec No C:1611–20.

66. Morris RJ, Fischer SM, Slaga TJ. Evidence that the centrally and peripherally located cells in the murine epidermal proliferative unit are two distinct cell populations. Journal of Investigative Dermatology. 1985;84:277–81.

67. Braun KM, Watt FM. Epidermal label-retaining cells: background and recent applications. J. Investig. Dermatol. Symp. Proc. 2004;9:196–201.

68. Zhang L, Li H, Zeng S, Chen L, Fang Z, Huang Q. Long-term tracing of the BrdU label-retaining cells in adult rat brain. Neurosci. Lett. 2015;591:30–4.

69. Nemeth K, Karpati S. Identifying the Stem Cell. Journal of Investigative Dermatology. Elsevier Masson SAS; 2014;134:1–5.

70. Ishizuya-Oka A. Epithelial-connective tissue cross-talk is essential for regeneration of intestinal epithelium. J Nippon Med Sch. 2005;72:13–8.

71. Lemons ML, Condic ML. Integrin signaling is integral to regeneration. Experimental Neurology. 2008;209:343–52.

72. Satoh A, Makanae A, Hirata A, Satou Y. Blastema induction in aneurogenic state and Prrx-1 regulation by MMPs and FGFs in Ambystoma mexicanum limb regeneration. Dev Biol. 2011;355:263–74.

73. Makanae A, Satoh A. Early regulation of axolotl limb regeneration. Anat Rec (Hoboken). Wiley Subscription Services, Inc., A Wiley Company; 2012;295:1566–74.

74. Monaghan JR, Athippozhy A, Seifert AW, Putta S, Stromberg AJ, Maden M, et al. Gene expression patterns specific to the regenerating limb of the Mexican axolotl. Biology Open. 2012;1:937–48.

75. Endo T, Bryant SV, Gardiner DM. A stepwise model system for limb regeneration. Dev Biol. 2004;270:135–45.

76. Wernig F, Mayr M, Xu Q. Mechanical stretch-induced apoptosis in smooth muscle cells is mediated by beta1-integrin signaling pathways. Hypertension. Lippincott Williams & Wilkins; 2003;41:903–11.

77. Fujisawa T. Role of interstitial cell migration in generating position-dependent patterns of nerve cell differentiation in Hydra. Dev Biol. 1989;133:77–82.

78. Boehm A-M, Bosch TCG. Migration of multipotent interstitial stem cells in Hydra. Zoology (Jena). 2012;115:275–82.

79. Mochizuki K, Sano H, Kobayashi S, Nishimiya-Fujisawa C, Fujisawa T. Expression and evolutionary conservation of nanos-related genes in Hydra. Dev Genes Evol. 2000;210:591–602.

80. Rebscher N, Volk C, Teo R, Plickert G. The germ plasm component Vasa allows tracing of the interstitial stem cells in the cnidarian Hydractinia echinata. Dev Dyn. Wiley**-**Liss, Inc; 2008;237:1736–45.

81. Gustafson EA, Wessel GM. Vasa genes: emerging roles in the germ line and in multipotent cells. Bioessays. WILEY**-**VCH Verlag; 2010;32:626–37.

82. Alié A, Leclère L, Jager M, Dayraud C, Chang P, Le Guyader H, et al. Somatic stem cells express Piwi and Vasa genes in an adult ctenophore: Ancient association of “germline genes” with stemness. Dev Biol. Elsevier Inc; 2011;350:183–97.

83. Plickert G, Frank U, Müller WA. Hydractinia, a pioneering model for stem cell biology and reprogramming somatic cells to pluripotency. Int J Dev Biol. 2012;56:519– 34.

84. Alié A, Hayashi T, Sugimura I, Manuel M, Sugano W, Mano A, et al. The ancestral gene repertoire of animal stem cells. Proc Natl Acad Sci USA. 2015;112:E7093–100.

85. Rinkevich B, Matranga V. Stem Cells in Marine Organisms. Springer Verlag; 2009.

86. Extavour CG, Pang K, Matus DQ, Martindale MQ. vasa and nanos expression patterns in a sea anemone and the evolution of bilaterian germ cell specification mechanisms. Evol Dev. 2005;7:201–15.

87. Bode HR. The interstitial cell lineage of hydra: a stem cell system that arose early in evolution. J Cell Sci. 1996.

88. Bosch TCG, Anton-Erxleben F, Hemmrich G, Khalturin K. The Hydra polyp: nothing but an active stem cell community. Dev Growth Differ. 2010;52:15–25.

89. Galliot B, Ghila L. Cell plasticity in homeostasis and regeneration. Mol. Reprod. Dev. Wiley Subscription Services, Inc., A Wiley Company; 2010;77:837–55.

90. Leclère L, Röttinger E. Diversity of cnidarian muscles: function, anatomy, development and regeneration. Front Cell Dev Biol. Frontiers; 2017;4:E3365.

